# Identification of a neural circuit that enables safe, long-term torpor in mice

**DOI:** 10.64898/2026.04.22.719582

**Authors:** Kexin Tong, Jingrui Yang, Feixiang Yuan, Min Tang, Yeting Gan, Shangming Wu, Xinyuan Tong, Peixiang Luo, Shanghai Chen, Hongbin Ji, Feifan Guo

## Abstract

Torpor, a state of regulated hypothermia and hypometabolism, is a critical survival strategy in certain mammals. Previous studies established the primacy of neuron control in initiating acute torpor. However, whether neuromodulation can safely sustain torpor over the long term, and the specific governing pathways, remain unanswered challenges limiting its translational potential. Here we identify a distinct preoptic neuronal population expressing General control nonderepressible 2 (‘G neurons’) in mice that is essential and sufficient for torpor induction. Strikingly, sustained and selective activation of G neurons induces a stable, weeks-long torpid state (G neuron-driven long-term torpor, ‘GLT’), from which animals arouse without obvious behavioral deficits or tissue pathology. In contrast, long-term torpor driven by pan-neuronal activation of the same brain region (PLT) causes multiple post-recovery damages, underscoring the unique safety profile of selective pathway of G neurons. In a mouse cancer model, GLT directly suppressed tumor proliferation and markedly sensitized tumors to chemotherapy, achieving profound therapeutic outcomes. Together, we delineate a distinct neuronal population safely enabling prolonged torpor. This establishes a specific paradigm for inducing a stable and sustained hypometabolic state, providing a transformative platform for fundamental physiological research and pioneering clinical strategies against chronic pathologies such as cancer.

## Introduction

To sustain homeostasis against environmental challenges, organisms have evolved complex physiological adaptations. Torpor, characterized by a profound reduction in body temperature and metabolic rate, represents an extreme but evolutionarily conserved energy-saving strategy to survive^1–4^. In situations where the energy is limited, rather than engaging in continuous, energetically expensive activities like foraging, torpor enables animals to dramatically reduce energy expenditure by periodically suppressing physiological functions, thereby enhances survival^1,5^. This phenomenon manifests in distinct patterns, ranging from multi-day bouts during hibernation to the daily torpor observed in non-hibernators such as laboratory mice, which display transient episodes of hypothermia and hypometabolism within a day^1,2^.

The neural architecture governing torpor is an area of intense investigation. Previous studies have implicated several key brain regions to be involved in torpor regulation, including the preoptic area (POA), dorsomedial hypothalamus, ventrolateral medulla, and the area postrema and nucleus of the solitary tract^6–11^. Within the POA, a central hub for thermoregulation^12,13^, specific neuronal populations have been identified as regulators for torpor induction, including those expressing pituitary adenylate cyclase-activating polypeptide (ADCYAP1)^10^, pyroglutamylated RFamide peptide (QRFP)^8^, estrogen receptor alpha (ERα)^9^, prostaglandin EP3 receptor (EP3R)^11^, and vesicular glutamate transporter 2 (VGLUT2)^7^. Nevertheless, this emerging neural diagram is incomplete, and other essential neuronal regulators remain to be discovered.

The prospect of safely inducing torpor has profound clinical implications^6^. The pursuit of therapeutically exploiting synthetic torpor has driven the development of various induction strategies, from systemic agents^6,14^ to modern neural approaches including chemogenetic/optogenetic manipulation^8–10^, targeted ultrasound^15^ and deep brain stimulation^7^. While these methods can elicit an acute torpor, holding clinical promise for acute scenarios like acute ischemia-reperfusion injury^7^ and organ transplantation^16^, their utility is constrained by a significant limitation: the induced state is invariably short-lived, lasting from hours to at most two days^8–10^. This temporal restriction is a major barrier to translating the benefits of hypothermia and hypometabolism to chronic diseases. This raises two crucial challenges in the field: first, whether a stable, long-term torpor can be achieved through certain means; second, whether animals can recover from such a prolonged state without incurring lasting physiological or behavioral damage, or if the specific biological pathway for its safe maintenance has simply remained elusive. Addressing these questions is essential for unlocking the full therapeutic potential of torpor.

Here, we turned our attention to General Control Nonderepressible 2 (GCN2), an evolutionarily ancient kinase that functions as a cellular sensor for amino acid deficiency^17^ and other nutritional stresses including fasting^18^, and mediates various adaptive changes crucial for survival. GCN2 is highly expressed in several brain regions^19–21^, including the amygdala, anterior piriform cortex, arcuate nucleus, and the medial POA (MPA)-a brain region known to be essential for orchestrating torpor in response to fasting^10^. These observations pinpointed GCN2-expressing neurons as a compelling, yet unexplored, candidate population that respond to fasting stimuli and play a regulatory role in the onset of torpor. Despite progress in understanding the diverse physiological functions of central GCN2, the roles of the neuronal population defined by its expression has never been addressed. Our study was undertaken to test the hypothesis and to explore the translational potential of this novel cell type.

Here, we report the discovery of a specific uncharacterized population of GCN2-expressing neurons within the anterior and ventral MPA (avMPA^GCN2^, designated as ‘G neurons’) that is essential and sufficient for torpor induction in mice. Remarkably, we establish that the selective, sustained activation of G neuron induce a stable, weeks-long, and fully reversible torpid state termed ‘GLT’ (G neuron-driven long-term torpor), and then leverage this synthetic state to reveal its therapeutic potential in a mouse cancer model.

## Results

### G neurons are activated during fasting-induced torpor and inhibition of these neurons abolishes fasting-induced torpor

To elucidate the neural mechanisms governing torpor induction, we established a fasting-induced daily torpor model, as previously described^10^, in 8-12-week-old male C57BL/6J wild-type (WT) mice (Fig. 1a). Upon 24-hour fasting, mice exhibited characteristic torpor bouts^10^, manifested as rapid decreases in core body temperature (T_core_) below 31 ℃^9,22^ with a sustained hypothermia followed by spontaneous arousal to baseline level, as recorded by implanted telemetric probes (Fig. 1b, c, Extended Data Fig. 1a). Infrared thermal imaging corroborated these observations, revealing synchronous changes during torpor in the surface temperature of brown adipose tissue (BAT) (T_BAT_), a major organ for non-shivering thermogenesis in mammals^23^, as well as rectal temperature (Extended Data Fig. 1b-d). The fasted mice further demonstrated profound hypometabolism, evidenced by markedly decreased metabolic rate parameters including oxygen consumption (VO_2_), energy expenditure (HEAT), and respiratory exchange ratio (RER, VCO_2_/ VO_2_) compared to fed controls (Fig. 1d, Extended Data Fig. 1e).

**Fig. 1.**
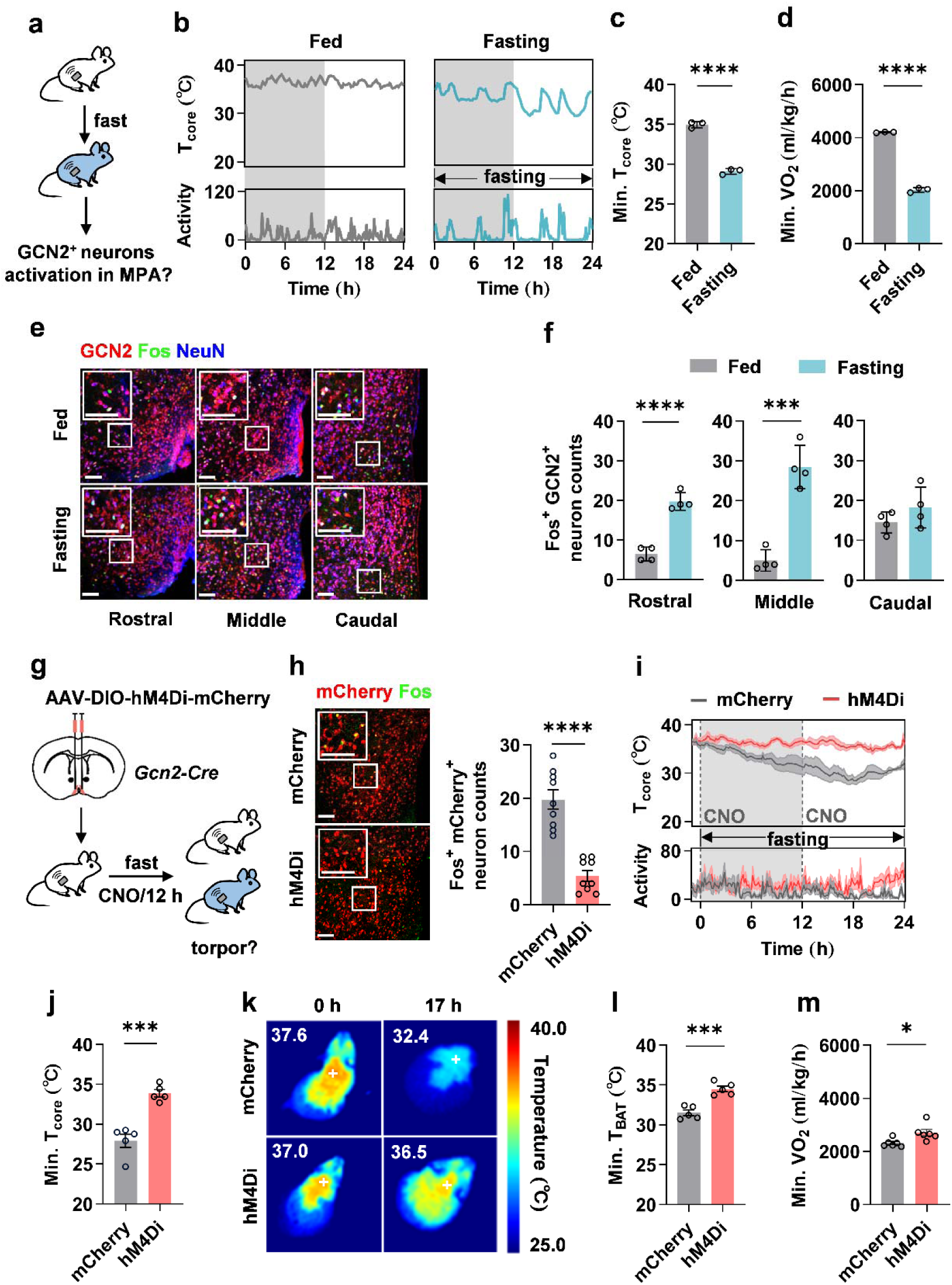
G neurons are activated by fasting and required for fasting-induced torpor. **a-f,** Wild-type mice were ad libitum fed (Fed) or fasted (Fasting) over 24 hours for neuronal activity examination. **a**, Experimental strategy. **b**, Representative traces of core body temperature (T_core_) and motor activity (Activity). Gray/white areas indicate 12 hours of darkness/light, respectively. **c, d,** Quantifications of the minimum T_core_ and metabolic rate (volume of oxygen consumed, VO_2_) (n = 3, ****p = 4.0e-5, ****p = 1.9e-6, respectively). **e,** Representative immunofluorescence (IF) staining images for GCN2 (red), Fos (green) and NeuN (blue) in rostral, middle or caudal medial preoptic area (MPA, bregma=0.50, 0.38 or -0.10) sections. Scale bars: 100 μm for both main panels and close-up. **f,** Quantifications of a Fos^+^/GCN2^+^/NeuN^+^ neuron counts in rostral, middle or caudal MPA (n = 4, ****p = 8.1e-5, ***p = 2.5e-4, p = 0.24, respectively). **g-m,** Chemogenetic inhibition of GCN2 expressing neurons within avMPA (G neurons) during fasting. **g,** Experimental strategy. **h,** Left, representative IF staining images for Fos (green) in avMPA sections in control (mCherry) and G-hM4Di (hM4Di) mice following CNO administration. Right, quantifications of Fos^+^/mCherry^+^ colocalized neuron counts (n = 9, ****p = 3.2e-6). Scale bars: 100 μm for both main panels and close-up. **i,** T_core_ and Activity before and during a 24-hour-fasting (n = 5). Dashed lines indicate the onset of CNO administration. **j,** Quantifications of minimum T_core_ (n = 5, ***p = 2.5e-4). **k,** Representative infrared thermal images of brown adipose tissue surface temperature (T_BAT_) at baseline (0 h) and after 17 h of fasting. **l, m,** Quantifications of minimum T_BAT_ and VO_2_ during fasting (n = 6, ***p = 3.3e-4, *p=0.019, respectively). Data are represented as mean ± SEM. Two-tailed unpaired Student’s t-test in (**c, d, f, h, i, k, l**). *p < 0.05; ***p < 0.001; ****p < 0.0001.

To investigate the involvement of GCN2-expressing neurons within MPA in fasting-induced torpor, we performed immunofluorescence (IF) analysis at 13 hours post-fasting, the time point corresponding to the steepest decline in body temperature observed in our and other’s study^24^. Triple IF staining for Fos (a neuronal activity marker^10^), GCN2, and NeuN (a neuronal marker^15^) revealed robust Fos expression in GCN2-expressing neurons specifically within avMPA (bregma - 0.50, 0.38 mm), but not in the posterior MPA (pMPA, bregma - 0.10 mm) (Fig. 1e, f). This anatomically restricted activation pattern was consistent with previous characterizations of MPA subregions in torpor regulation^8,10^.

To determine whether GCN2-expressing neurons within avMPA are functionally required for torpor, we employed chemogenetic manipulation using our newly generated *Gcn2-Cre* transgenic line that enables Cre-dependent targeting of GCN2-expressing cells (Extended Data Fig. 2a). Adeno-associated virus (AAV) encoding Cre-dependent inhibitory Designer Receptors Exclusively Activated by Designer Drugs (DREADDs) (AAV-DIO-hM4Di-mCherry) were bilaterally injected into avMPA in *Gcn2-Cre* mice, while AAV encoding only mCherry (AAV-DIO-mCherry) were injected as controls (Fig. 1g). Systemic administration of clozapine-*N*-oxide (CNO), a DREADDs agonist^10^ effectively suppressed Fos expression in hM4Di-expressing neurons (Fig. 1h), confirming the functional inhibition. Critically, CNO-mediated inhibition of G neurons during fasting prevented the development of hypothermia (T_core_ below 31 ℃) and T_BAT_ reduction characteristic in fasting-induced torpor (Fig. 1i-l, Extended Data Fig. 2b). Additionally, minimum oxygen consumption increased in inhibition group compared to controls (Fig. 1m). Under fed state, no significant differences in thermoregulatory or metabolic parameters were observed between two groups, except for transiently elevated T_BAT_ and rectal temperatures immediately following CNO injection in inhibited group (Extended Data Fig. 2c-h). Together, these results demonstrated that the neuronal activity of G neurons, marked by GCN2 within avMPA, is essential for fasting-induced torpor.

To verify whether G neurons comprise a functionally distinct population, we characterized their neuronal composition and relationship to previously identified torpor-regulating neuronal populations in MPA^9,10^. GCN2 and NeuN co-staining revealed that G neurons constitute approximately 68.0% of all neurons, evenly distributed throughout the region (Extended Data Fig. 3a). To determine the neurochemical identity of these neurons, we performed GCN2 and NeuN co-staining in transgenic mice expressing fluorescent reporters under control of *Vglut2* (excitatory glutamatergic neuron marker^8^) or vesicular GABA transporter (*Vgat*, inhibitory GABAergic neuron marker^8^) promoters (*Vglut2*-ai9, *Vgat*-ai9 mice). Among G neurons, 27.75% were excitatory (*Vglut2*^+^) and 53.07% were inhibitory (*Vgat*^+^) (Extended Data Fig. 3b, c), indicating a heterogeneous population. Furthermore, 32.75 % of G neurons co-expressed ERα, and 42.92 % co-expressed ADCYAP1 (Extended Data Fig. 3d, e). While G neurons partially overlap with previously characterized torpor-regulating neurons, the large substantial proportion of non-overlapping confirmed that G neurons functions as a previously unidentified neuronal population regulating torpor.

### Activation of G neurons alone is sufficient to promote torpor in mice

To examine whether activation of G neurons is sufficient to trigger torpid state, we bilaterally injected *Gcn2-Cre* mice with AAV encoding Cre-dependent excitatory DREADDs (AAV-DIO-hM3Dq-mCherry) into avMPA (G-hM3Dq mice), while AAV encoding only mCherry (AAV-DIO-mCherry) were injected as control group (Fig. 2a). Activation of hM3Dq-expressing G neurons following CNO administration was verified by Fos staining (Fig. 2b). Following CNO administration, G-hM3Dq mice exhibited a rapid drop in T_core_ that fell below 31 °C and persisted for more than 72 h, whereas control mice maintained T_core_ above ∼35 °C throughout the period (Fig. 2c), with T_BAT_ declined in a parallel (Fig.2d, e). Additionally, a sustained and significant hypometabolism was observed in G-hM3Dq mice compared to control group (Fig. 2f, Extended Data Fig. 4a, b). G-hM3Dq mice also presented a similar rectal temperature changes, a slight decrease in body weight, with a significant reduction in food intake (Extended Data Fig. 4c-e). Comparable torpor phenotypes were observed in female G-hM3Dq mice (Extended Data Fig. 5a-g). To further validate torpor induction effects in an independent methodology, we activated G neurons optogenetically in mice receiving AAV encoding a Cre-dependent excitatory ChR2 (AAV-DIO-ChR2-mCherry)^8^, and observed similar torpor phenotypes on thermoregulation compared to control group (Extended Data Fig. 4f-k). Together, our results demonstrated that targeted activation of G neurons is sufficient to promote a profound torpor.

**Fig. 2.**
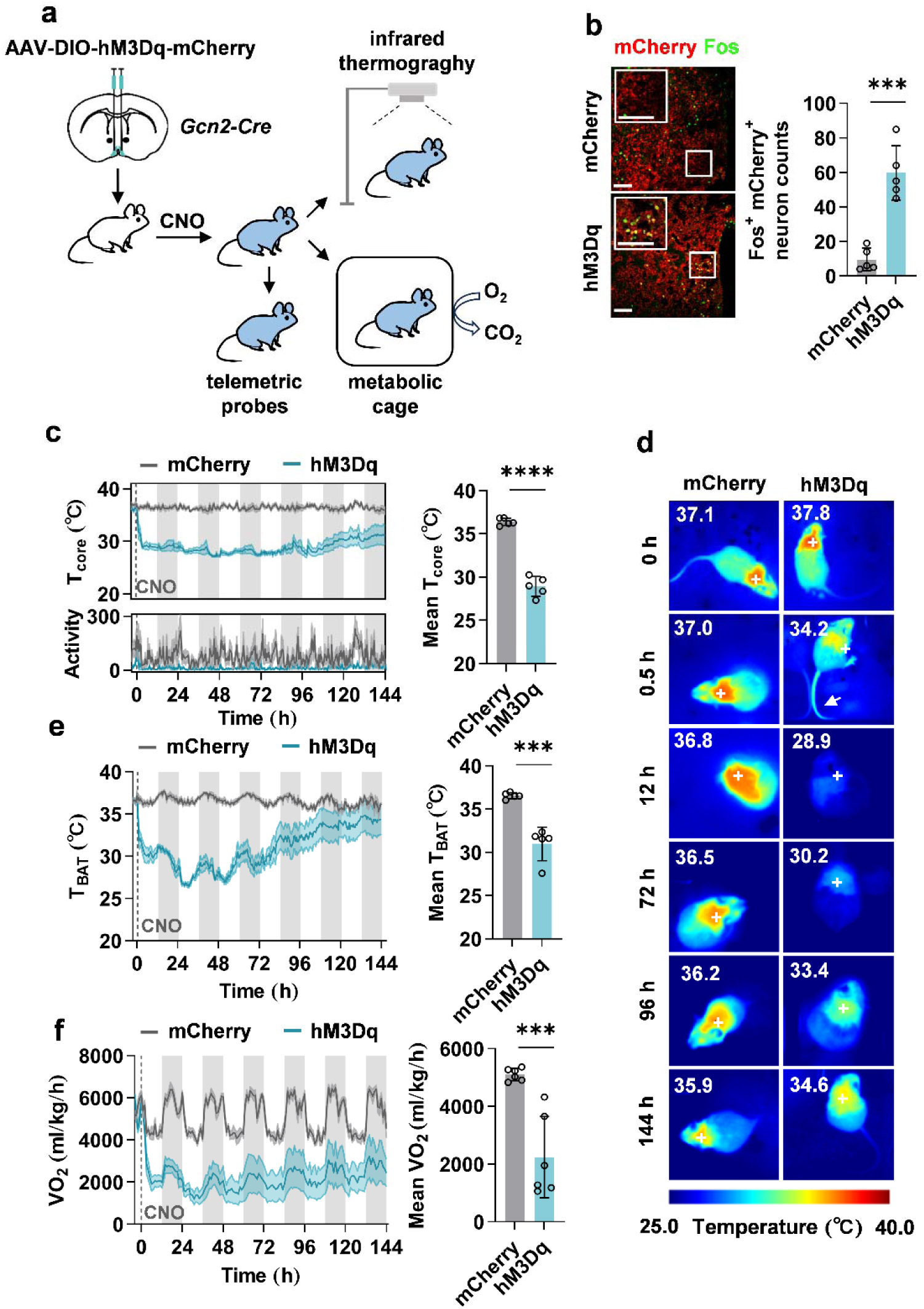
Activation of G neurons alone is sufficient to promote torpor in mice. **a**, **Experimental** strategy. **b,** Left, representative immunofluorescence (IF) staining images for Fos (green) in avMPA sections in control (mCherry) and G-hM3Dq (hM3Dq) mice following CNO administration. Right, quantifications of Fos^+^/mCherry^+^ colocalized neuron counts (n = 5, ***p = 1.7e-4). Scale bars: 100 μm for both main panels and close-up. **c,** Left, core body temperature (T_core_) and motor activity (Activity) before and after CNO administration (n = 5). Right, quantifications of minimum T_core_ post-injection (n = 5, ****p = 7.9e-7). Gray/white areas indicate 12 hours of darkness/light, respectively. Dashed lines indicate the onset of CNO administration. **d,** Representative infrared thermal images of brown adipose tissue surface temperature (T_BAT_) at time indicated before and after CNO administration. Note the transient increase in tail temperature (arrow at 0.5 h), indicative of cutaneous vasodilation. **e,** Left, T_BAT_ before and after CNO administration. Right, quantifications of mean T_BAT_ after CNO administration (n = 5, ***p = 2.3e-4). **f,** Left, metabolic rate (volume of oxygen consumed, VO_2_) before and after CNO administration. Right, quantifications of mean VO_2_ after CNO administration (n = 6, ***p = 5.9e-4). Data are represented as mean ± SEM. Two-tailed unpaired Student’s t test in (**b, c, e, f**). ***p < 0.001; ****p < 0.0001.

### Repeated activation of G neurons induces a long-term torpor

Given that G neuron-driven torpor following single administration of CNO could last up to 72 h, and caused no observable damages, we explored whether sustained activation of these neurons could extend acute torpor into a sustained torpid state with prolonged time scale. We administered CNO every 48 h over 24 d, a protocol designed to provide repeated neuronal stimulation (Fig. 3a). G-hM3Dq mice presented maintained suppressed rectal temperature throughout the entire experimental period compared with controls, with no evidence of spontaneous arousal or temperature recovery (Fig. 3b). Consistently, other parameters including T_BAT_ and metabolic rate parameters (VO, HEAT, and RER) remained significantly suppressed at both mid-term (d 12-14) and final-term (d 24-26) measurement intervals (Fig. 3c–f; Extended Data Fig. 6a-d). As a critical control, repeated CNO administrations alone in WT mice produced no thermoregulatory changes (Extended Data Fig. 7a-c), confirming that the sustained torpor phenotype is attributable specifically to hM3Dq-mediated activation of G neurons.

**Fig. 3.**
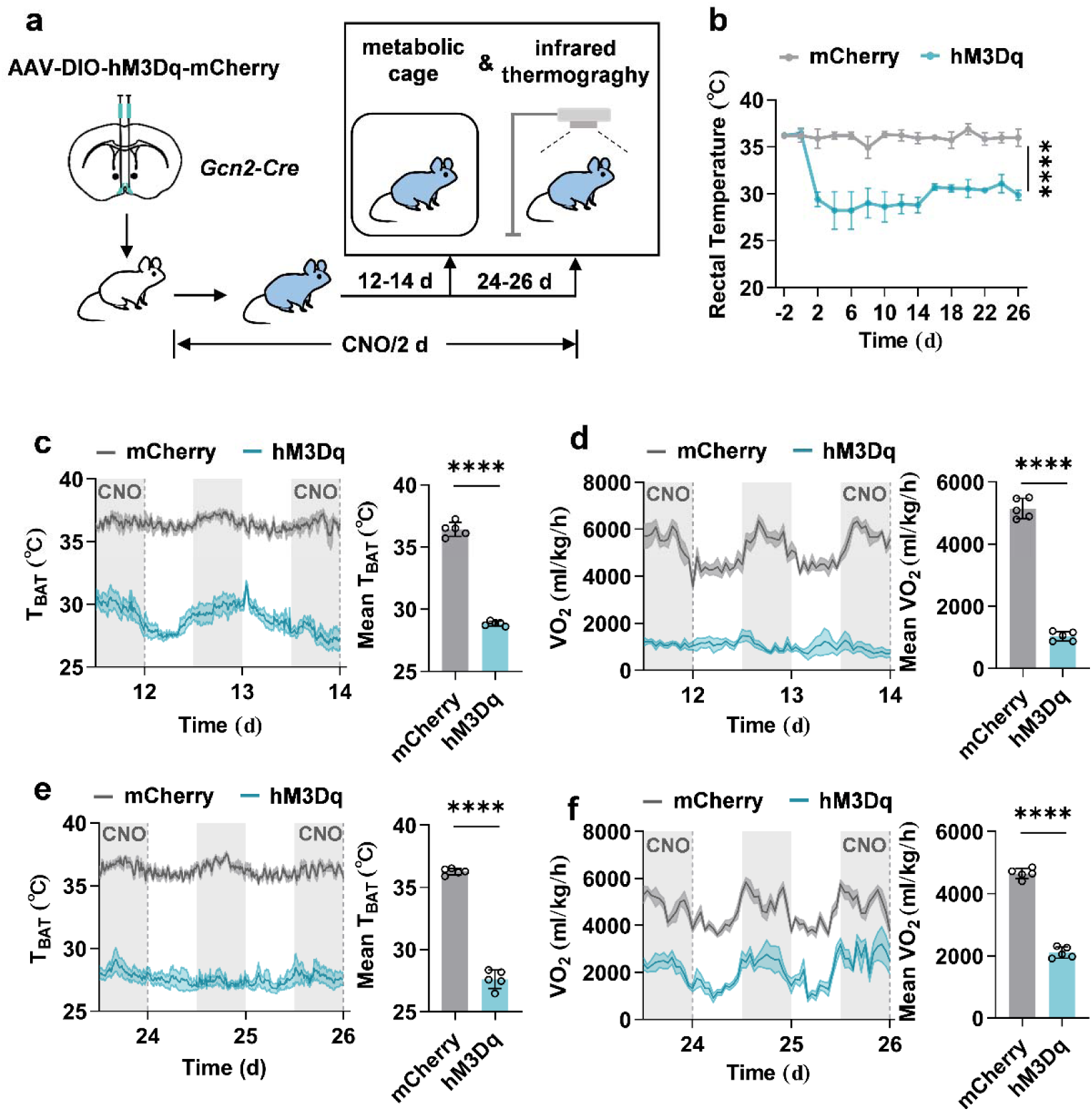
Repeated activation of G neurons induces a G neuron-driven long-term torpo (GLT). **a,** Experimental strategy for GLT induction and phenotypes measurement. **b,** Rectal temperature of control (mCherry) and G-hM3Dq (hM3Dq) mice at indicated time (n = 5, ****p = 2e-15). **c, d,** Left, brown adipose tissue surface temperature (T_BAT_) and the metabolic rat of oxygen consumed, VO_2_) during mid-term (12-14 day). Right, quantifications of the mean and the mean VO_2_ (n = 5, ****p = 3.0e-9, ****p = 5.8e-9, respectively). Gray/white areas indicat hours of darkness/light, respectively. Dashed lines indicate the onset of CNO administration. **e, f,** Left, T_BAT_ and VO_2_ during final term (24-26 day). Right, the quantifications of the mean T_BAT_ VO_2_ (n = 5, ****p = 7.5e-9, ****p = 1.1e-8, respectively). Data are represented as mean ± SEM. Two-tailed unpaired Student’s t test in (**c, d, e, f**). RM ANOVA with Geisser-Greenhouse’s correction in (**b**). ****p < 0.0001.

To address whether reduced food intake during long-term G neurons activation might account for the observed sustained torpor, we subjected WT mice to 50% caloric restriction, which produced food intake levels comparable to those of G-hM3Dq mice in repeated activation (Extended Data Fig. 6f, 7d). Though 50% caloric restriction induced a transient mild decrease in rectal temperature during the initial three days, it failed to establish or sustain a stable long-term hypothermic state (Extended Data Fig. 7f), accompanied by severe weight loss which not present in G-hM3Dq mice (Extended Data Fig. 6e, 7e). This marked divergence in thermogenic responses between the neuronal activation *versus* metabolic restriction demonstrated that sustained, stable long-term torpor required direct activation of G neurons, and could not be achieved through caloric restriction alone. These findings showed that repeated chemogenetic activation of G neurons was sufficient to induce a previously unreported G neuron-driven long-term torpor, termed GLT, maintaining throughout the entire process without evidence of recovery or compensatory arousal.

### Behavioral competence and tissue integrity are preserved following GLT

To evaluate the safety profile of prolonged GLT, we conducted comprehensive behavioral and physiological assessments in G-hM3Dq mice following the 26-day GLT induction protocol (Fig. 4a). We first assessed whether GLT produces any behavioral deficits. A series of standardized behavior tests^8^, including open filed test (OFT), elevated plus maze (EPM), tail suspension test (TST), and rotarod test (RR) were performed after GLT termination (Fig. 4a). Across all four behavioral tests, G-hM3Dq mice exhibited indistinguishable performance compared to control group (Fig. 4b-e), demonstrating preserved exploratory behavior, anxiety-related measures, depressive-like behavior, and motor coordination. Moreover, daily monitoring revealed that body weight, food intake, and rectal temperature of G-hM3Dq mice returned to baseline levels within the recovery period, comparable to control group (Extended Data Fig. 8a-c). Consistently, tissue integrity and morphology were similarly preserved. Comprehensive histological analysis for fat tissues (subcutaneous white adipose tissue, sWAT; epididymal white adipose tissue, eWAT; BAT), liver, skeletal muscle, heart, kidney, and brain revealed no differences in morphology, with no differences in tissue weight compared to control group (Fig. 4f, Extended Data Fig. 8d). Together, these observations demonstrated that GLT is physiologically tolerable, with no detectable behavior deficits or tissues damages.

**Fig. 4.**
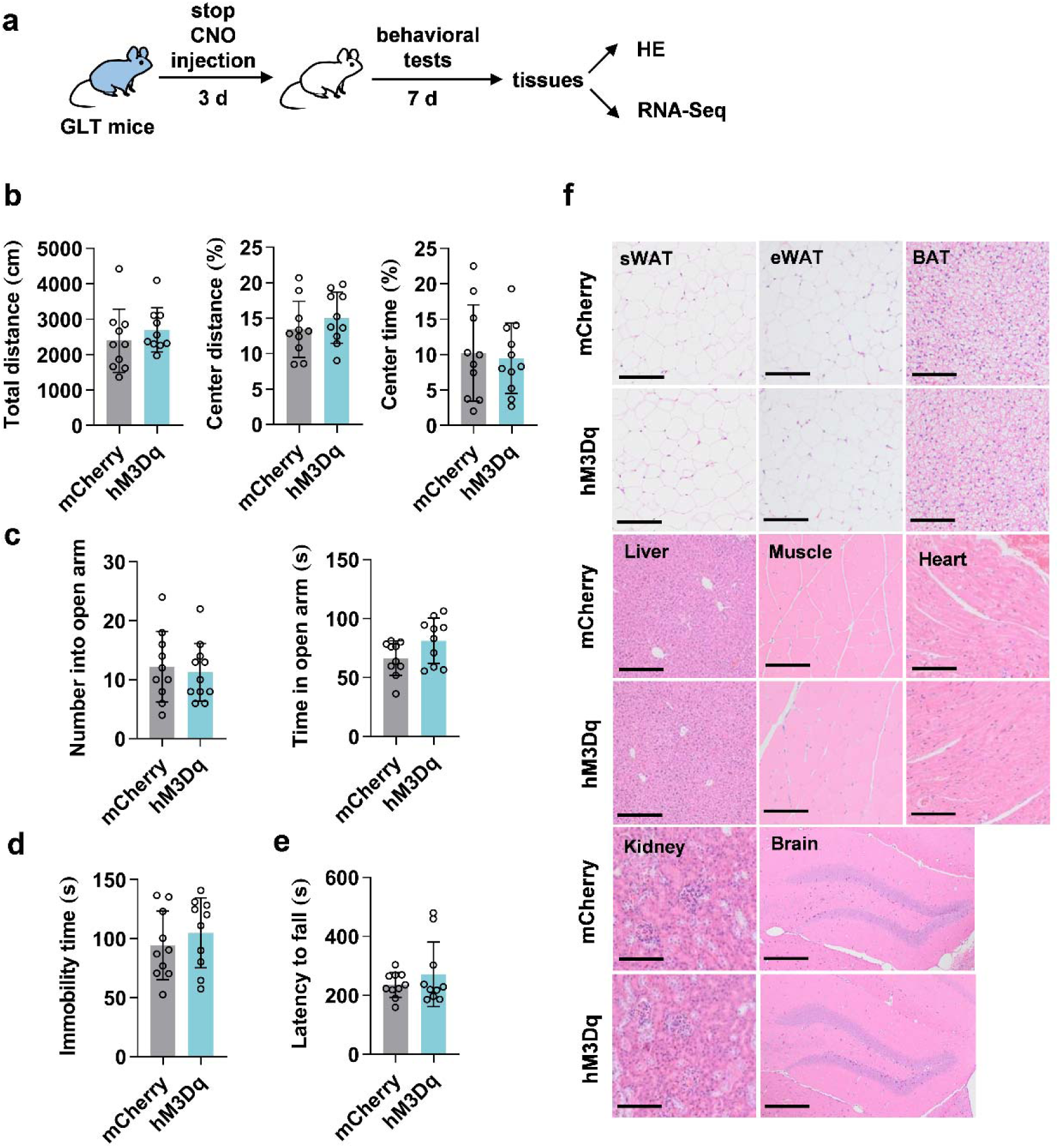
No secondary pathological changes are observed following G neuron-driven long-term torpor (GLT). **a,** Experimental strategy for behavior tests and tissue analysis following GLT. Open filed test (OFT), elevated plus maze (EPM), tail suspension test (TST), and rotarod test (RR), hematoxylin and eosin staining (HE). **b,** Total travel distance (Total distance), percentage of distance in center area (Central distance), and percentage of time spent in center (Central time) of OFT following the induction in control (mCherry) and G-hM3Dq mice (hM3Dq, n = 10, p = 0.39, p = 0.35, p = 0.77, respectively). **c,** Number of entries into th open arms (Number into open arm) and time spent in the open arms (Time in open arm) of EPM (n = 10, p = 0.70, p = 0.069, respectively). **d,** Immobility time of TST (n = 10, p = 0.43). **e,** Latency to fall of RR (n = 10, p = 0.35). **f,** Representative images of HE staining of subcutaneous white adipose tissue (sWAT), epididymal white adipose tissue (eWAT), brown adipose tissue (BAT), liver, muscle, heart, kidney and brain. Scale bars: 200 μm for sWAT, eWAT, BAT, muscle, heart, kidney and brain; 400 μm for liver. Data are represented as mean ± SEM. Two-tailed unpaired Student’s t test in (**b, c, d, e**).

To provide a more comprehensive understanding of the safety profile of GLT, we performed a direct comparison with torpor induced by pan-neurons of avMPA. Previous studies demonstrated that acute torpor can be elicited by activating all avMPA neurons^10,25^, however, whether repeated pan-neuronal activation of avMPA could induce long-term torpor remained unknown. Moreover, the relative safety profiles of selective (G neurons) *versus* non-selective (pan-neurons) activation for torpor induction had not been systematically compared. To address these questions, we bilaterally injected AAV encoding hM3Dq pan-neuronally under the *hSyn* promoter^25^ (AAV-hSyn-hM3Dq-mCherry) into avMPA of WT mice, while AAV only encoding mCherry (AAV-hSyn-mCherry) were injected as control (Extended Data Fig. 9a). Single CNO administration in avMPA-hM3Dq mice produced similar acute torpor phenotypes reported by others^25^ which extended over 48 hours (Extended Data Fig. 9b, c). Moreover, repeated CNO administration (every 48 h for 24 d) similarly induced a hypothermic and hypometabolic pan-neuron-driven long-term torpor (PLT) (Extended Data Fig. 9a, d-g). However, unlike G-hM3Dq mice, avMPA-hM3Dq mice exhibited multiple post-recovery deficits: despite the rectal temperature and food intake returned to baseline levels after torpor termination, body weight declined progressively and failed to recover (Extended Data Fig. 9h-j); behavioral tests revealed anxiety-like phenotypes (reduced center-zone time and distance proportion in OFT, decreased open-arm exploration in EPM and increased tendency in immobility time in TST) and impaired motor function (reduced movement in OFT and decreased latency to fall in RR; Extended Data Fig. 9l-o). Consistent with body weight and behavior tests, recovered avMPA-hM3Dq mice displayed lower eWAT and gastrocnemius muscle mass and histological evidence of lipid-droplet reduction and myofibrillar atrophy, respectively (Extended Data Fig. 9k, p).

To evaluate systemic tissue safety at the molecular level, we performed RNA sequencing on key tissues (BAT, eWAT, liver and skeletal muscle) collected after GLT and PLT (Fig. 4a, Extended Data Fig. 9a). In PLT-treated mice, we observed marked transcriptional dysregulation characterized by the hyperactivation of pathways and marker genes related to inflammation, cell death/stress, and tissue fibrosis, with the eWAT and liver being the most severely affected (Extended Data Fig. 10a, b), corroborated the macroscopic tissue atrophy and behavioral deficits observed in the PLT group. Conversely, GLT intervention elicited no such pathological transcriptomic responses. The expression profiles of the same functional domains in GLT mice remained highly stable across all tissues examined (Extended Data Fig. 10a, b).

These results established that GLT was a safe, well-tolerated physiological state with preservation of behavioral competence, tissue integrity, and molecular homeostasis. In contrast, pan-neuronal activation of avMPA triggered secondary pathological changes across multiple organ systems and behavioral domains. In conclusion, our comprehensive analysis highlighted the unique safety profile of GLT.

### GLT suppresses tumor growth and enhances chemotherapeutic efficacy

To examined whether GLT could function as a chronic diseases management, such as a direct anti-tumor intervention, we utilized primary lung adenocarcinoma cells derived from *Kras^G12D/+^*; *Trp53^fl/fl^* mice (KP cells)^26^ to establish a tumor model in G-hM3Dq mice. Beginning at d 6 post-implantation, when baseline tumor volumes were comparable between groups, GLT induction was initiated via repeated CNO administration every 48 h (Fig. 5a). Surprisingly, the induction of GLT potently restricted tumor progression, whereas control group exhibited rapid tumor expansion, reaching ethical limits by d 24 post-implantation (Fig. 5b). The volumes and weights of tumors at the study endpoint were significantly lower in GLT-treated group (Fig. 5b-d). Histological analysis of resected tumors corroborated the macroscopic findings, revealing a stark morphological shift from the hypercellular, solid tumor architecture characteristic of rapid proliferation in control group to a markedly reduced cellular density with disrupted tissue organization in the GLT-treated group (Fig. 5e). These findings demonstrated that GLT was sufficient to suppress tumor growth.

**Fig. 5.**
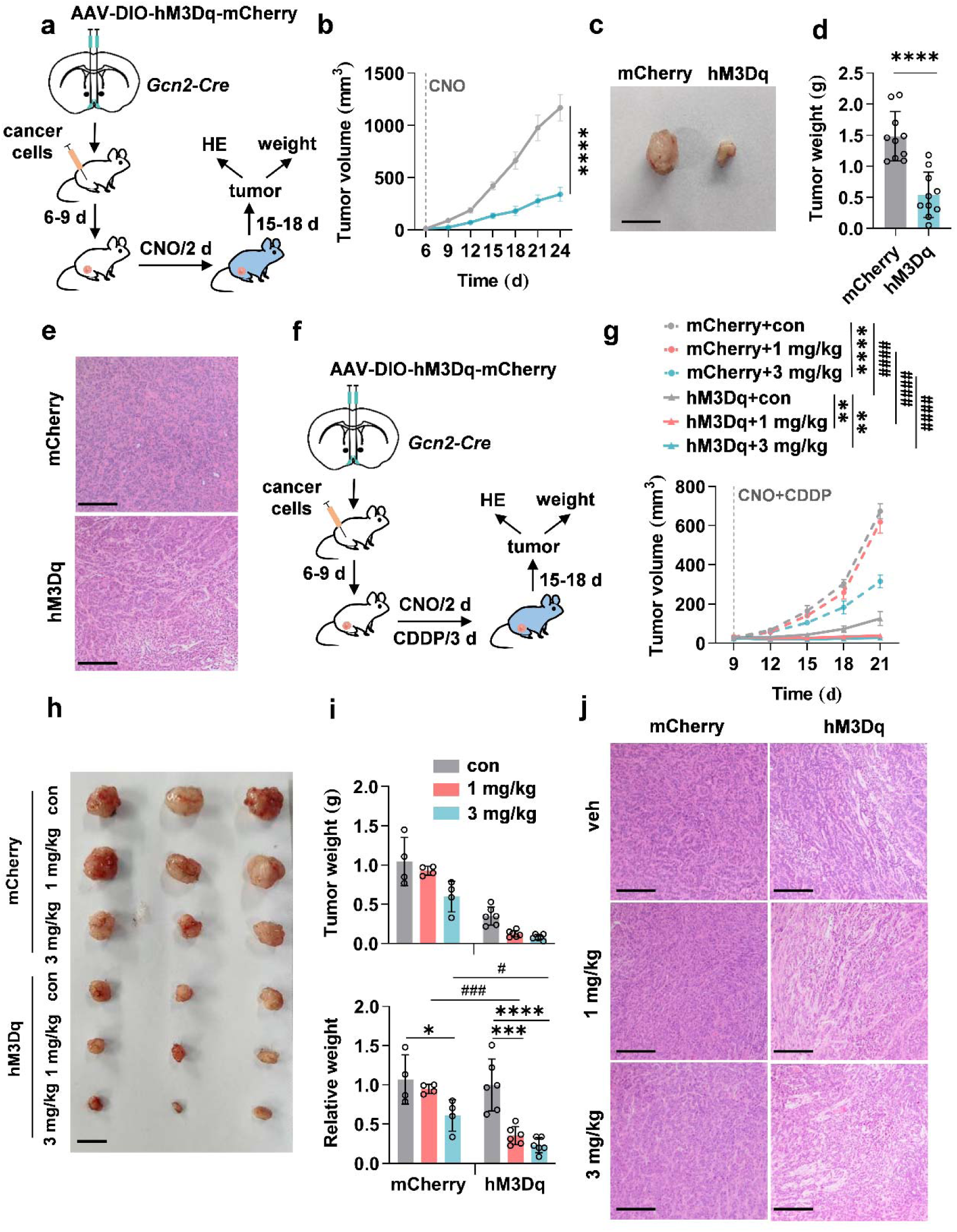
G neuron-driven long-term torpor (GLT) impairs tumor growth and enhances chemotherapy. **a-e,** Direct tumor suppression effects of GLT. **a,** Experimental strategy, hematoxylin and eosin staining (HE). **b,** Tumor volume growth from d 6-24 after implantation in control (mCherry, n = 14) and G-hM3Dq mice (hM3Dq, n = 13, ****p = 6e-15). Dashed lin indicates the beginning of CNO administration. **c, d, e,** Representative images, weight, hematoxylin and eosin (HE) staining images of tumors harvested on d 24 after implantation (n = 10, ****p = 3.2e-5). Scale bars: 1cm for (**c, h**), 500 μm for (**e, j**). **f-j,** Effects of GLT in chemotherapy. **f,** Experimental strategy. **g,** Tumor volume growth from d 9-21 after implantation in non-GLT group with different drug treatments (vehicle, 1mg/kg cisplatin [cis-Diaminodichloroplatinum, CDDP] and 3mg/kg CDDP) (mCherry+con, n = 5, mCherry+1mg/kg, n = 5, mCherry+3mg/kg, n = 6) and GLT group (hM3Dq+con, hM3Dq+1mg/kg, hM3Dq+3mg/kg, n = 8) (mCherry+con *vs* mCherry+1mg/kg, p = 0.70, mCherry+Con *vs* mCherry+3mg/kg, ****p = 1.2e-11, hM3Dq+con *vs* hM3Dq+1mg/kg, **p = 5.4e-3, hM3Dq+con *vs* hM3Dq+3mg/kg, **p = 5.5e-3, mCherry+con *vs* hM3Dq+con, ^####^p < 1e-15, hM3Dq+1mg/kg *vs* hM3Dq+1mg/kg, ^####^p < 1e-15, hM3Dq+3mg/kg *vs* hM3Dq+3mg/kg, ^####^p < 1e-15). Dashed line indicates the beginning of drugs administration. **h, i, j,** Representative images, weight, HE staining images of tumors harvested on d 21 after implantation. **i,** Up, tumor weight. Below, tumor weight respectively relative to control in mCherry and hM3Dq groups (mCherry, n = 4, hM3Dq, n = 6, mCherry+con *vs* mCherry+1mg/kg, p = 0.72, mCherry+con *vs* mCherry+3mg/kg, *p = 0.037, hM3Dq+con *vs* hM3Dq+1mg/kg, ***p = 2.4e-4, hM3Dq+con *vs* hM3Dq+3mg/kg, ****p = 3.9e-5, hM3Dq+1mg/kg *vs* hM3Dq+1mg/kg, ^###^p = 7.1e-4, hM3Dq+3mg/kg *vs* hM3Dq+3mg/kg, ^#^p = 0.033). Data are represented as mean ± SEM. Two-tailed unpaired Student’s t test in (**d**). Two-way RM ANOVA with Geisser-Greenhouse’s correction in (**b,g**). One-way ANOVA with Tukey’s multiple comparisons test and two-way ANOVA with Geisser-Greenhouse’s correction followed by post hoc unpaired t-test with Bonferroni’s correction in (**i**). ^*/#^p < 0.05; **p < 0.01; ^***/###^p < 0.001; ^****/####^p < 0.0001.

We next investigated whether GLT could enhance the anti-tumor efficacy of chemotherapy. We combined GLT induction with administration of cisplatin (cis-Diaminodichloroplatinum, CDDP)^27^, a widely used chemotherapy agent, in tumor-bearing G-hM3Dq mice (Fig. 5f). We first examined the anti-tumor efficacy of cisplatin monotherapy across two dose levels. Low-dose cisplatin alone showed negligible effects on tumor growth, whereas high-dose cisplatin produced partial tumor suppression (Fig. 5g-j). We next administered these two cisplatin doses with concurrent GLT induction. As expected, GLT alone recapitulated the potent tumor suppression. Surprisingly, when combined with chemotherapy, GLT broadened the therapeutic window. The previously ineffective low-dose cisplatin achieved significant anti-tumor effects compared to vehicle-treated group under GLT, which was absent in non-GLT mice (Fig. 5g-j). Additionally, GLT also augmented the anti-tumor efficacy of high-dose cisplatin, as reflected by a greater reduction in relative tumor weight under GLT induction, compared with non-GLT mice (Fig. 5i, below). Histological analysis of tumor also demonstrated markedly reduced cellular proliferation in mice receiving GLT induction combined with low-dose cisplatin, consistent with the volumetric findings (Fig. 5j).

In conclusion, these findings suggested that GLT both independently suppressed tumor growth and synergized with chemotherapy to achieve enhanced anti-tumor effect.

## Discussion

While the central mechanisms governing acute torpor entry are increasingly well characterized^10^, the neural architecture required to maintain a prolonged, damage-free hypometabolic state has remained largely uncharted. In this work, we identify G neurons, a distinct population of preoptic neurons marked by GCN2 that are both essential for daily torpor and sufficient to initiate synthetic torpor in mice. Repeated activation of G neurons enables the induction of GLT, a stable and harmless long-term torpor (Extended Data Fig. 11). This discovery provides a powerful new tool for dissecting the neurobiology of torpor, and establishes a dedicated paradigm for inducing a long-term hypothermic and hypometabolic state with significant therapeutic potential, which we demonstrate in a mouse cancer model.

While the functions of nutrient sensing and metabolism control of the GCN2 kinase in the CNS are known^19,28^, the physiological role of GCN2-expressing neurons remains completely unknown. Here, we observed that G neurons, as a previously uncharacterized GCN2 expressing neuronal population in avMPA, become robustly active during fasting-induced torpor, indicating that these neurons may play roles in torpor. To interrogate their function, we generated a novel *Gcn2-Cre* mouse line. Using this tool, we demonstrated that chemogenetic silencing of G neurons abrogates the hypothermic and hypometabolic response to fasting, confirming their necessity for daily torpor. These neurons represent a population largely distinct from previously described torpor-regulating neurons (ADCYAP1, ERα, or VGLUT2 expressing neurons)^7,9,10^, establishing G neurons as a novel neuronal component of the torpor regulatory network.

Beyond their role in daily torpor, chemogenetic activation of G neurons was sufficient to induce a profound torpor in both male and female mice that persisted over 72 hours, a duration notably longer than those elicited by stimulating other known torpor-regulating neurons (QRFP^8^, ADCYAP1^10^, ERα^9^, EP3R^11^ or VGLUT2^7^ expressing neurons, lasting hours to 48 h). This superior effect suggests a unique and potent role inherent to this population, and may stem from its molecular or anatomical profile. We speculate that the prolonged duration of torpor may be attributed to the broader and more central neural innervation, as non-colocalized subset of G neurons may command distinct downstream circuits or deploy unique molecular effectors inaccessible to other torpor-regulating populations. Dissecting the specific contributions of these subsets will be a vital future endeavor to understand how G neurons orchestrate a physiological program of such remarkable duration and stability.

A crucial challenge in the field has been discovering how to extend hypothermia and hypometabolism beyond the acute phase^6^. This limitation likely stems from the transient nature of conventional neuromodulatory approaches, which often fail to sustain physiological suppression. Our study provides a breakthrough in this regard. The profound efficacy of G neurons suggested that their manipulation could bypass these constraints, facilitating the transition from acute to prolonged torpor via repeated, low-frequency interventions. By implementing a regimen of repeated CNO administration every two days, we successfully induced GLT, a stable G neuron-driven long-term torpor. The measurements of thermal and metabolic state at different intervals confirmed persistent, profound hypothermia and hypometabolism throughout the entire protocol, with no phenotypic attenuation or habituation observed at the final stage compared to the middle one. This further demonstrated the stability of GLT, and the obvious feasibility to extending its duration. Correctively, we establish a new and robust paradigm for inducing long-term torpid state, moving far beyond the inherent limitations of acute models.

Given the profound physiological shifts inherent to torpor, inquiry into how to safely induce and sustain a long-term torpid state has remained the ultimate prerequisite for clinical translation. Our comprehensive evaluation following GLT arousal revealed no evidence of behavioral deficits or histological damage, with mice fully maintaining normal body and organ weights. To further establish the unique safety profile of GLT, we performed a comparative analysis by pioneering an equally novel paradigm of long-term torpor driven by broad pan-neuronal activation of all avMPA neurons termed PLT. While PLT successfully sustained prolonged long-term torpor with basic physiological functions remained intact, it triggered certain notable post-recovery detriments, including specific behavioral alterations and histological abnormalities in muscle and eWAT. Consistently, RNA-Seq analysis revealed a broad transcriptomic footprint of systemic damages in PLT group, which was absent in GLT group. These observations highlight that while broad avMPA activation can drive a similar long-term torpor, the subpopulation of G neurons constitute a highly refined and specific locus for safe regulation. The adverse off-target effects of PLT likely stem from the concurrent activation of non-G neurons, recruiting distinct, deleterious downstream circuits not engaged by G neurons. Ultimately, isolating this specific G neuron pathway resolves the critical challenge of safely inducing and sustaining long-term torpor, minimizing systemic disruption and unlocking its true therapeutic potential.

The distinct safety and precision of GLT open new therapeutic avenues, particularly for chronic diseases. While acute torpor has been proposed for acute injuries^6^, whether the long-term torpor could be utilized for specific diseases remains unknown. A recent study shows torpor might slow blood epigenetic aging, suggesting the possibility for its application in the management of chronic diseases such as cancer. Under prolonged continuous hypometabolic conditions, cellular proliferation was significantly suppressed^3^, offering a potential approach for inhibiting tumor growth, given that the core characteristic of tumor cells is uncontrollable proliferation^29^. Here, we provide the first direct evidence that GLT can, by itself significantly suppress tumor growth. Furthermore, combining GLT with two different doses of cisplatin resulted in a synergistic anti-tumor effect that was more effective than either monotherapy. This approach not only enhances chemotherapy efficacy but also holds the promise of mitigating dose-dependent side effects^27^. While our work demonstrated that the tumor suppression arises from the systemic hypothermic and hypometabolic state, the further mechanism requires future determination. Here, we combined GLT with chemotherapy, which remarkably suppressed the tumor growth, introducing a fundamentally new strategy for cancer treatment.

In summary, our work identified G neurons, a previously uncharacterized neuronal population, as a key regulator of safely sustaining long-term torpor, thereby establishing a novel and robust methodology to achieve this elusive state in non-hibernators. This breakthrough work opens several compelling avenues for future investigation. First, the precise upstream signals activating G neurons during fasting, and the downstream circuits orchestrating torpor, remain to be fully elucidated. Given that GCN2 is a well-established nutrient-stress sensor mediating metabolic responses in both peripheral and central systems^19,28,30–32^, we hypothesize that GCN2 in avMPA acts as a central transducer, translating fasting-induced nutrient scarcity into the neuronal activation required for torpor. Second, while our current data highlight the unique safety profile of GLT, comprehensive long-term profiling is necessary to assess any potential subtle or delayed systemic effects. Third, the high evolutionary conservation of GCN2 suggests that targeting homologous neuronal populations may induce a similar hypothermic and hypometabolic state in other species. Future studies in larger mammals, particularly non-human primates, will be an essential step toward evaluating translational feasibility and safety. Finally, unraveling the precise molecular mechanisms by which GLT suppresses tumor growth and enhances chemotherapy could unveil novel biological targets. Beyond oncology, this underscores the broader potential applicability of GLT to other chronic diseases and pathological states.

Addressing these questions will be critical in translating this safe, sustained hypothermic and hypometabolic model into transformative therapeutic strategies.

## Methods

### Animals

All animal experiments were conducted in accordance with the guidelines of the Institutional Animal Care and Use Committee at Fudan University (No. 2022030006S) and the Institutional Animal Care and Use Committee of Shanghai Institute of Biochemistry and Cell Biology, Chinese Academy of Sciences (2023-037). All mice were on a the C57BL/6J genetic background. Wild-type (WT) mice were obtained from the Model Animal Research Center of Nanjing University (Nanjing, China). The *Gcn2*-P2A-Cre mouse line was generated by CRISPR/Cas9-mediated homologous recombination in C57BL/6J embryonic stem cells (see Extended Data Fig. 1). *Vglut2-*IRES-Cre mice (Stock No. 016963), *Vgat-*IRES-Cre (Stock No. 016962), and Ai9 (tdTomato) reporter mice (Stock No. 007909) (Bar Harbor, ME, USA). To visualize glutaminergic and GABAergic neurons, *Vglut2*-IRES-Cre mice and *Vgat*-IRES-Cre mice were respectively intercrossed with Ai9 reporter mice. All experiments were performed using male mice unless otherwise indicated.

Mice were housed under a 12:12 h light/dark cycle (lights on at 08:00) at an ambient temperature of 22-23°C, with ad libitum access to water and rodent standard chow diet (Shanghai Pu Lu Teng Biotechnology, P1103F). For caloric restriction experiments, mice were provided 50 % of their average daily food intake by weight. Mice were sacrificed by CO_2_ inhalation.

### Fasting-induced daily torpor inductions

Adult mice (8-12 weeks old) were single-housed overnight for adaption before induction. The food is removed at the beginning of the dark cycle (20:00)^10^ at an ambient temperature of 22-23 °C. Torpor was defined when the core temperature was equal to or below 31 °C^24^. Torpor entries of mice were observed about 8-12 h after fasting. After 24 h of fasting, the mice were given food again.

### Long-term torpor inductions

During the long-term torpor induction, mice were singly housed. CNO injection were performed every 48 h for 24 d to maintain torpid state. Food intake, rectal temperature and body weight were recorded every 48 h for 24 d. Serum glucose level, body composition, BAT temperature and indirect calorimetry were measured at mid-term (d 12-14) and the final term (d 24-26). Mice then were allowed 3 d for recovering before behavioral tests. After the behavioral tests, mice were sacrificed and the tissues were harvested.

### Subcutaneous tumorigenesis and chemotherapy treatments

All experiments involving subcutaneous tumorigenesis employed primary lung adenocarcinoma cells derived from *Kras^G12D/+^*; *Trp53^fl/fl^* mice (KP cells)^26^. All experiments with subcutaneously inoculated KP cells were performed using 7-8-week-old WT or *GCN2-Cre* mice. 5 × 10^6^ Kras/p53 cells suspended in 100 μl of Matrigel (#354234, Corning, Shanghai, USA) were injected subcutaneously into the dorsal flank^33^. The injection site was shaved and disinfected prior to tumor cell implantation. Nine days post-injection, mice with palpable tumors were randomized into different groups. Tumor growth was monitored every three days by measuring the longest (L) and shortest (S) diameters with a Vernier caliper, and tumor volume was calculated as (L × S²)/2. Mice were euthanized when tumor size reached the humane endpoint in our home office license (2000 mm^3^ in volume or 15 mm in any direction).

In chemotherapy treatments, Cisplatin (MedChem Express, NJ, USA) was administered i.p. at 1 or 3 mg/kg, prepared from a 10 mg/ml stock in N,N-Dimethylformamide (DMF) (MedChem Express, NJ, USA) diluted in saline. Control mice received a vehicle of saline with an equivalent concentration of DMF at a dose of 3 mg/kg.

### Stereotaxic surgeries

Stereotaxic surgery was performed using a stereotaxic frame (RWD Life Science, Shenzhen, China). Mice body temperature was maintained using a heating pad throughout the surgery. Ophthalmic ointment was applied to maintain eye lubrication. Viruses were injected at a rate of 50 nL/min using a micro syringe pump connected to glass pipettes. Viruses were bilaterally injected into the avMPA (coordinates from bregma: AP +0.45 mm, ML ± 0.25 mm, DV -5.2 mm from bregma). After injection, the glass pipettes were left in place for 8 min before withdrawal to allow for diffusion. In mice involving optogenetic manipulations, optical fibres were then implanted unilaterally above the avMPA (AP +0.45 mm, ML 0.0 mm, DV -5.15 mm). The mice were allowed to recover from anesthesia on a heat blanket and were then intraperitoneally injected with antibiotics (ceftriaxone sodium, 0.1 g/kg) for 3 d to prevent infection. Mice were allowed 3-4 weeks for recovery and viral expression. Data were only included if these viruses were targeted specifically to G neurons or the fibre-optic implants were precisely placed.

### DREADDs

Neuronal populations were selectively activated or inhibited using Cre-dependent AAVs in *Gcn2-Cre* mice or a pan-neuronal AAV in WT mice. Viruses were bilaterally injected into the avMPA (150 nL per hemisphere).

#### G neurons Activation

A Cre-dependent AAV encoding an excitatory DREADD GPCR (AAV9-EF1a-DIO-hM3Dq-mCherry, 2.5 × 10^12^ genome copies per mL) at a volume of 150 nL was stereotaxically injected, or an AAV encoding only mCherry (AAV9-EF1a-DIO- mCherry, 2.5 × 10^12^ genome copies per mL) as control. CNO was administered i.p. at 0.2 mg/kg.

#### G neurons Inhibition

A Cre-dependent AAV encoding an inhibitory DREADD GPCR (AAV9-EF1a-DIO-hM4Di-mCherry, 3.5 × 10^12^ genome copies per mL) at a volume of 150 nL was stereotaxically injected, or an AAV encoding only mCherry (AAV9-EF1a-DIOmCherry, 3.5 × 10^12^ genome copies per mL) as control. CNO was administered i.p. at 0.4 mg/kg.

#### Pan-neuronal Activation within avMPA

A neuron-dependent AAV encoding an excitatory DREADD GPCR (hM3Dq) (AAV9-hSyn-hM3Dq-mCherry, 2.5 × 10^12^ genome copies per mL) at a volume of 150 nL was stereotaxically injected, or an AAV encoding only mCherry (AAV9-hSyn-mCherry, 2.5 × 10^12^ genome copies per mL) as control. CNO was administered i.p. at 0.2 mg/kg.

### Optogenetics manipulation

To activate G neurons, a Cre-dependent AAV encoding an ChR2 (AAV9-EF1a-DIO-ChR2-mCherry, 3 × 10^12^ genome copies per mL) at a volume of 150 nL was stereotaxically injected, or an AAV encoding only mCherry (AAV9-EF1a-DIO- mCherry, 2.5 × 10^12^ genome copies per mL) as control.

Mice were adapted for 1 h after attaching implanted cannulated fibre to optical fibre patch cable using ceramic sleeves (200-μm diameter, Thinkertech, NanJing, China). Patch cables were connected to 473nm lasers (473-nm blue, Thinkertech, NanJing, China) for applying optogenetic manipulations. The laser power at the optic fibre tip was adjusted to 8–10 mW, and the laser stimulation was applied at 20 Hz with 1 s ON/1 s OFF duty cycle for 30 min.

### Telemetric monitoring of core body temperature and gross motor activity

Mice were singly housed and implanted abdominally with telemetric probes (TA-F10, Data Sciences International, MN, USA). After at least seven days of recovery, mice were recorded in standard cages placed onto a radio frequency receiver platform (Data Sciences International, MN, USA). Core body temperature and gross motor activity were logged every 60 seconds.

### Metabolic parameters measurements

BAT surface temperature was measured by an infrared camera (Magnity Electronics Co., Ltd., Shanghai, China). At least 3 d prior to the experiment, the mice were anesthetized, and the hair on their dorsal scapular area was carefully shaved off using a razor to fully expose the skin covering BAT. This allowed infrared camera to accurately capture the local surface temperature of the BAT. The rectal temperature of mice was measured using a digital thermometer (Physitemp Instruments, Clifton, NJ, USA). Indirect calorimetry was measured by metabolic cage (CLAMS-16; Columbus Instruments, USA). Mice were maintained in comprehensive lab animal monitoring system with food and water for 24-60 h. The dynamic parameters of indirect calorimetry were continuously recorded. The body composition of mice was detected by a Bruker Minispec mq10 NMR Analyzer (Bruker, Billerica, MA, USA).

### Behavioral tests

All behavioral tests were performed in the afternoon after a 1-hour acclimation period. Arenas were cleaned with 75% ethanol between trials.

#### OFT

Mice were placed in the center of a white plastic open field arena (50 cm × 50 cm × 50 cm) and allowed to explore freely for 10 min. A video camera positioned directly above the arena was used to track animal movements, recorded on a computer with LabState (AniLab) to determine the total distance and the time spent in the center of the chamber compared to the edges.

#### EPM

The EPM consisted of a central platform (5 × 5 cm^2^), two closed arms with walls, and two opposing open arms without walls (25 cm × 5 cm). The maze was placed 60 cm above the floor. Mice were placed on the central platform facing an open arm and was allowed to explore the maze for 5 min. The time spent in the open arms and the number of entries into the open arms, were analyzed using LabState (AniLab).

#### RR

Rotarod tests were performed using an accelerating rotarod (Ugo Basile) in which mice were placed on the rotating drum (5.7 cm diameter). The initial speed of the rotarod was set at 5 rpm, and the speed gradually increased from 5 to 60 rpm within 600 s. A trial ended if the mouse fell off the drum or the gripped the device and spun around for 2 consecutive revolutions. The latency time of the mice was recorded.

### Immunofluorescence (IF) staining

Mice were transcardially perfused with saline followed by 4% paraformaldehyde (PFA). Brains were dissected and post-fixed overnight in 4% PFA, followed by cryoprotection in PBS containing 20 % and 30 % sucrose. Free-floating coronal sections (25 μm) were prepared with a cryostat. Slices were blocked for 1 h at room temperature in PBST (0.3 % Triton X-100) with 5 % normal donkey serum, followed by incubation with primary antibodies at 4 °C overnight and secondary antibodies at room temperature for 2 h.

Primary antibodies included rabbit anti-GCN2 (1:500, Abcam, Cambridge, UK); rat anti-Fos (1:1000, Asis Biofarm, ZheJiang, China); rabbit anti-NeuN (1:1000, Proteintech, IL, USA), mouse anti-ERα (1:500, Santa Cruz Biotechnology, CA, USA), mouse anti-adcyap1 (1:500, Santa Cruz Biotechnology, CA, USA).

### Histological Analysis

Liver, BAT, eWAT, sWAT, heart, kidney, muscle, brain and tumor were fixed in 4 % PFA for 14-16 h. The tissues were then embedded in paraffin and the entire prepared block was cut into 8-µm sections. After deparaffinization and rehydration, the sections were stained with hematoxylin and eosin (H&E).

### RNA Sequency analysis

Total RNA was extracted from tissues using TRIzol (Invitrogen, Carlsbad, California, USA) according to manual instruction. Total RNA was qualified and quantified using a Fragment Analyzer or Agilent 2100 Bioanalyzer (Agilent, CA, USA), or Qseq-400 (Bioptic, Taiwan, China). Library preparation is performed using Optimal Dual-mode mRNA Library Prep Kit (BGI-Shenzhen, China). The final DNA nanoballs (DNBs) were sequenced on the G400/T7/T10 platform (BGI-Shenzhen, China), generating PE 100/150 reads.

The sequencing data was filtered, and the clean reads were mapped to the reference genome. Expression level of gene was calculated, and the differential expression analysis was performed using the DESeq2 (v1.34.0)^34^ with an adjusted P-value ≤ 0.05.

To take insight to the change of phenotype, KEGG (https://www.kegg.jp/) enrichment analysis of annotated different expression gene was performed by Phyper based on Hypergeometric test. All bioinformatic analyses were conducted using the Dr. Tom Multi-omics Data Mining System (BGI).

### Statistical analysis and reproducibility

Statistical analyses were performed in GraphPad Prism 8.0 (GraphPad Software, San Diego, CA). All values are presented as the mean ± standard error of the mean (SEM). When two groups were compared, data were analyzed for statistical significance either using a two-tailed unpaired Student’s t test or a two-way RM analysis of variance (ANOVA) with Geisser-Greenhouse’s correction followed by post hoc unpaired t-test with Bonferroni’s correction as indicated in the figure legends. For experiments involving multiple comparisons, data were analyzed for statistical significance using a one-way ANOVA with Tukey’s multiple comparisons test. For transparency, individual data points are shown on all relevant graphs. P values are indicated within the graphs. Statistical significance was set at p < 0.05.

## Author contributions

K.T., J.Y. and F.G. designed the study; F.G., F.Y. acquired the funding; K.T., J.Y., F.Y., M.T., Y.G., S.W., P.L., and S.C. performed animal experiments; X.T., and H.J. provided the KP cell line, K.T., J.Y. and F.G. performed the statistical analyses; F.G., K.T. and J.Y. wrote the manuscript.

## Data, code, and materials availability

All data are available in the main text or the supplementary materials.

## Extended Figures and Figure legends

**Extended Data Fig. 1.**
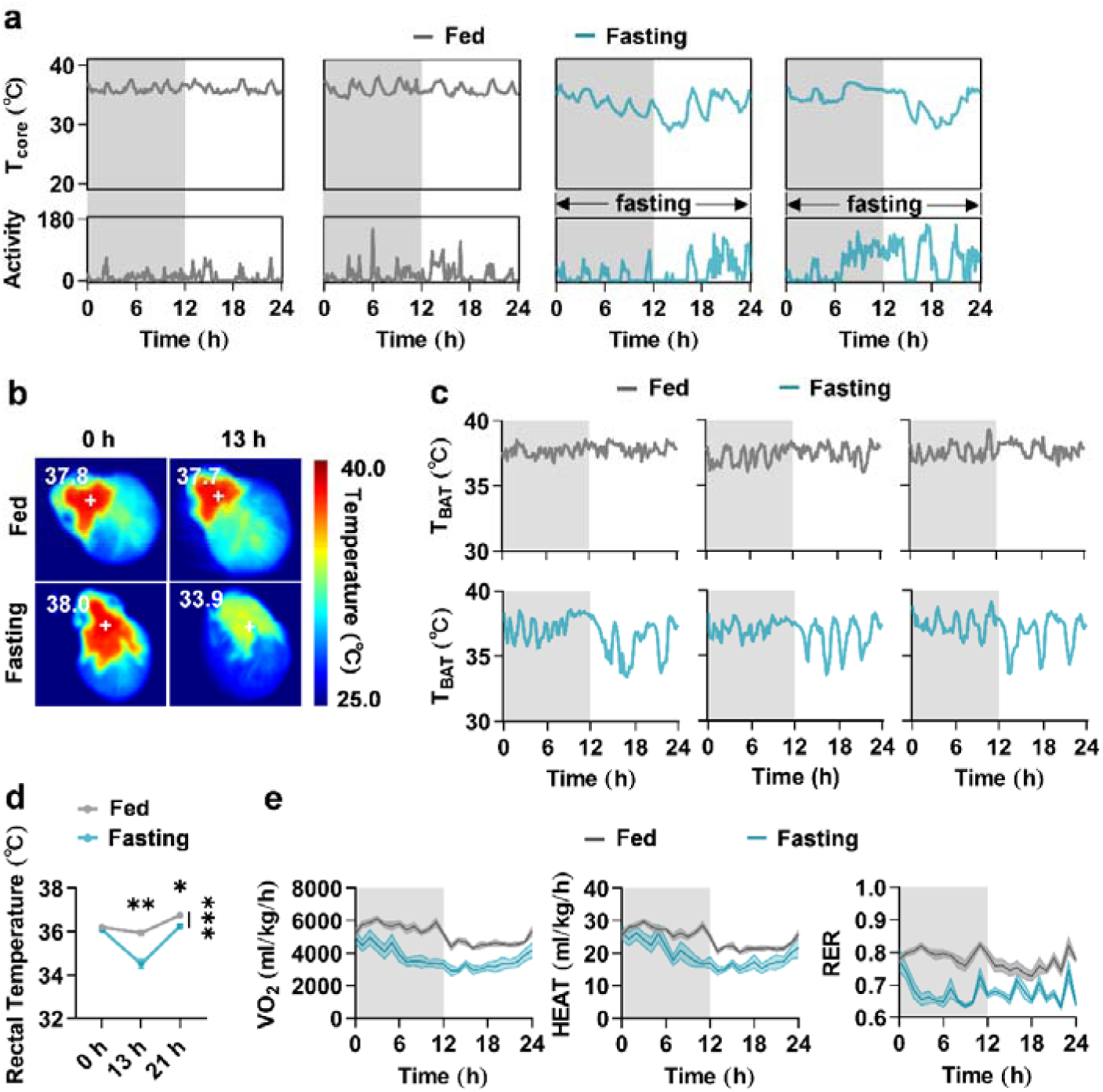
Body temperature and metabolic rate decrease during fasting-induced torpor. **a,** Traces of core body temperature (T_core_) and motor activity (Activity) in ad libitum fed (Fed) and fasted (Fasting) wild-type mice over 24 hours. Gray/white indicate 12 hours of darkness/light, respectively. **b,** Representative infrared thermal images of brown adipose tissue surface temperature (T_BAT_) at baseline (0 h) and 13 h. **c,** Traces of T_BAT_ during 24 h. **d,** Rectal temperature at baseline (0 h), 13 h, 21 h (n = 6, ***p = 2.7e-4 between lines, p = 0.41, **p = 1.3e-3, *p = 0.012, at time indicated respectively). **e,** Metabolic rate (volume of oxygen consumed, VO_2_), energy expenditure (HEAT), and respiratory exchange ratio (RER) during 24 h (n = 8). Data are represented as mean ± SEM. Two-way RM ANOVA with Geisser-Greenhouse’s correction followed by post hoc unpaired t-test with Bonferroni’s correction in (**d)**. *p < 0.05; **p < 0.01; ***p < 0.001.

**Extended Data Fig. 2.**
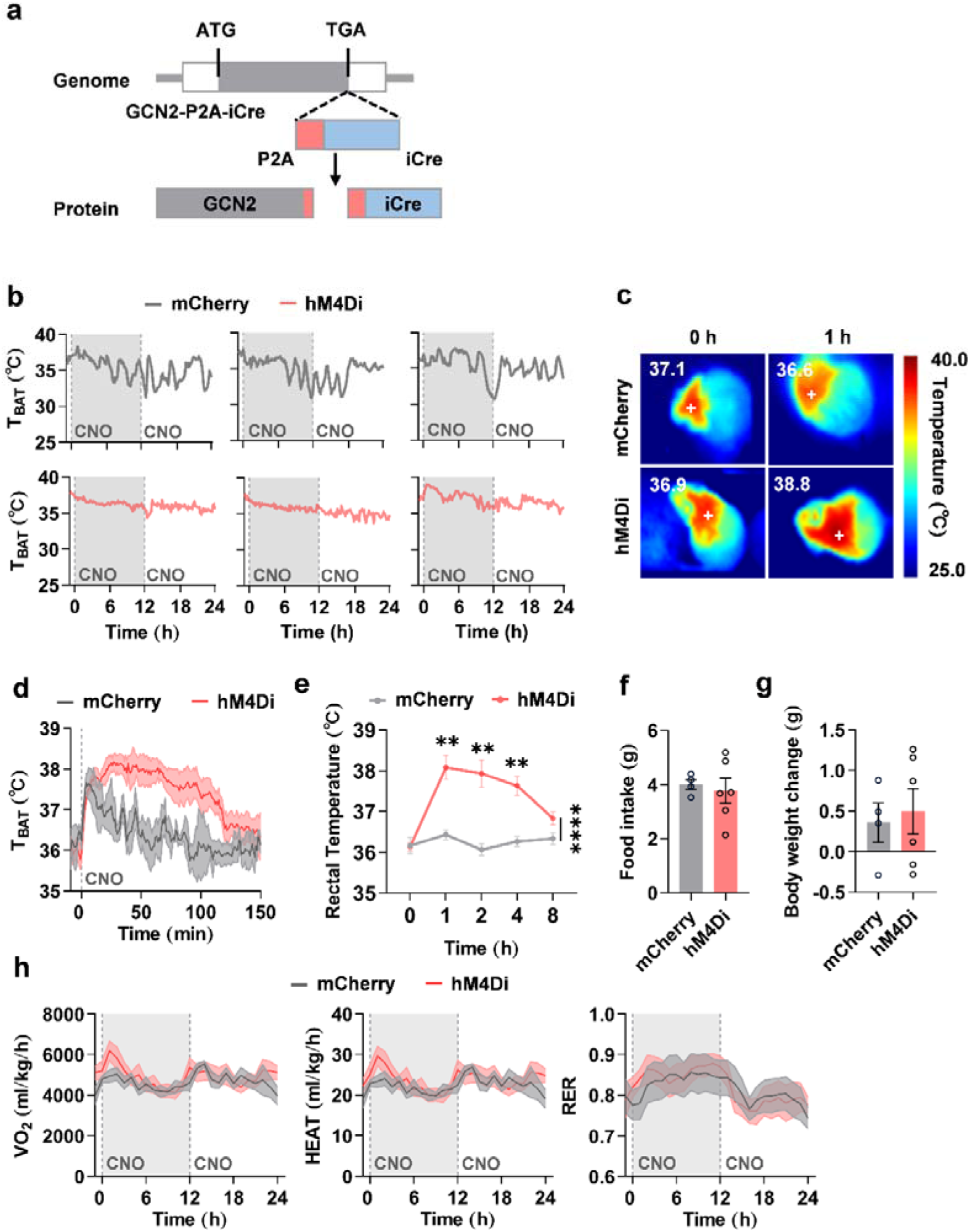
Generation of *Gcn2-Cre* mouse line, effects of inhibition of G neurons under fed and fasted state. **a,** Schematic of the *Gcn2^P2A-iCre^* locus. **b,** Traces of brown adipose tissue surface temperature (T_BAT_) before and during a 24-hour-fasting in control (mCherry) mice and G-hM4Di (hM4Di) mice. Gray/white indicate 12 hours of darkness/light, respectively. Dashed lines indicate the onset of CNO administration. **c-h,** The effects of inhibition of G neurons under fed state. **c,** Representative infrared thermal images of T_BAT_ at baseline (0 h) and 1 h following CNO administration. **d,** Traces of T_BAT_ before and after CNO administration (mCherry, n = 3, hM4Di, n = 6). **e,** Rectal temperature at indicated time (mCherry, n = 3, hM4Di, n = 6, p = 1, **p = 7.9e-3, **p = 7.9e-3, **p = 7.0e-3, p = 0.29, respectively). **f, g,** Total food intake and body weight change over 24 h (mCherry, n = 4, hM4Di, n = 6, p = 0.71, p = 0.74, respectively). **h,** Metabolic rate (volume of oxygen consumed, VO_2_), energy expenditure (HEAT), and respiratory exchange ratio (RER) (n = 5). Data are represented as mean ± SEM. Two-way RM ANOVA with Geisser-Greenhouse’s correction followed by post hoc unpaired t-test with Bonferroni’s correction in (**d**). Two-tailed unpaired Student’s t test in (**e, f, g**). **p < 0.01.

**Extended Data Fig. 3.**
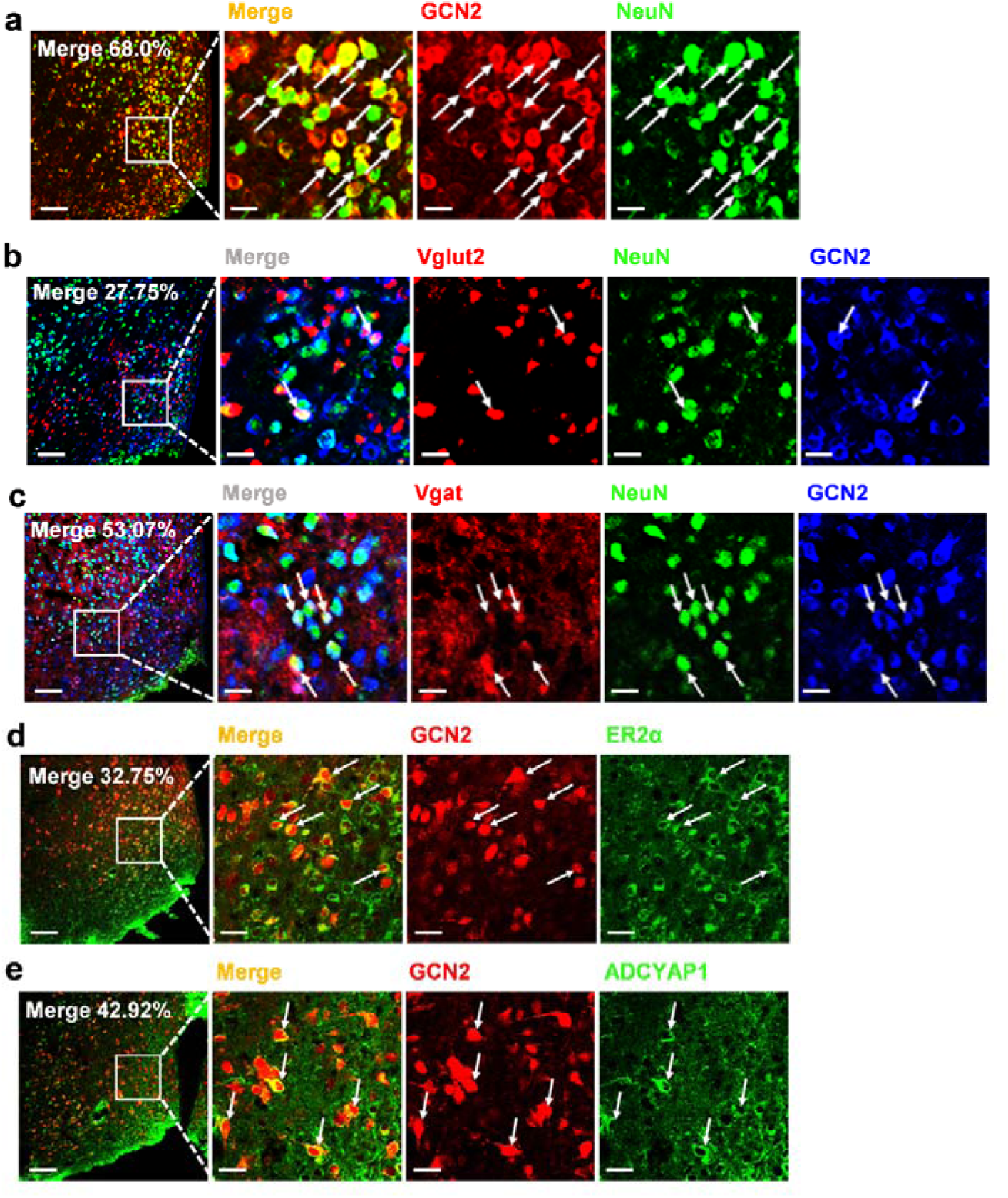
G neurons represent a novel population distinct from other torpor-regulating populations. **a,** Representative immunofluorescence (IF) staining images for NeuN (green) and GCN2 (blue) and the percentage of G neurons in total neurons (n=8). Scale bars: 100 μm for main panels and 25 μm for close-up. **b, c,** *Vglut2*-Ai9 (**b**) or *Vgat*-Ai9 (**c**) mice were utilized to discussed the proportion of excitatory or inhibitory neurons in G neurons. Representative IF staining images for NeuN (green) and GCN2 (blue) and the percentage of *Vglut2*^+^ or *Vgat*^+^ neurons in G neurons (n=10, n=8 respectively). **d, e,** G-mCherry mice were utilized to discussed the proportion of ER2α^+^ or ADCYAP1^+^ neurons in G neurons. Representative IF staining images for ER2α (from 14 mice) (**d**) or Adcyap1 (from 12 mice) and the percentage of ER2α^+^ or ADCYAP1^+^ neurons in G neurons (n=14, n=12 respectively). (**e**) (green).

**Extended Data Fig. 4.**
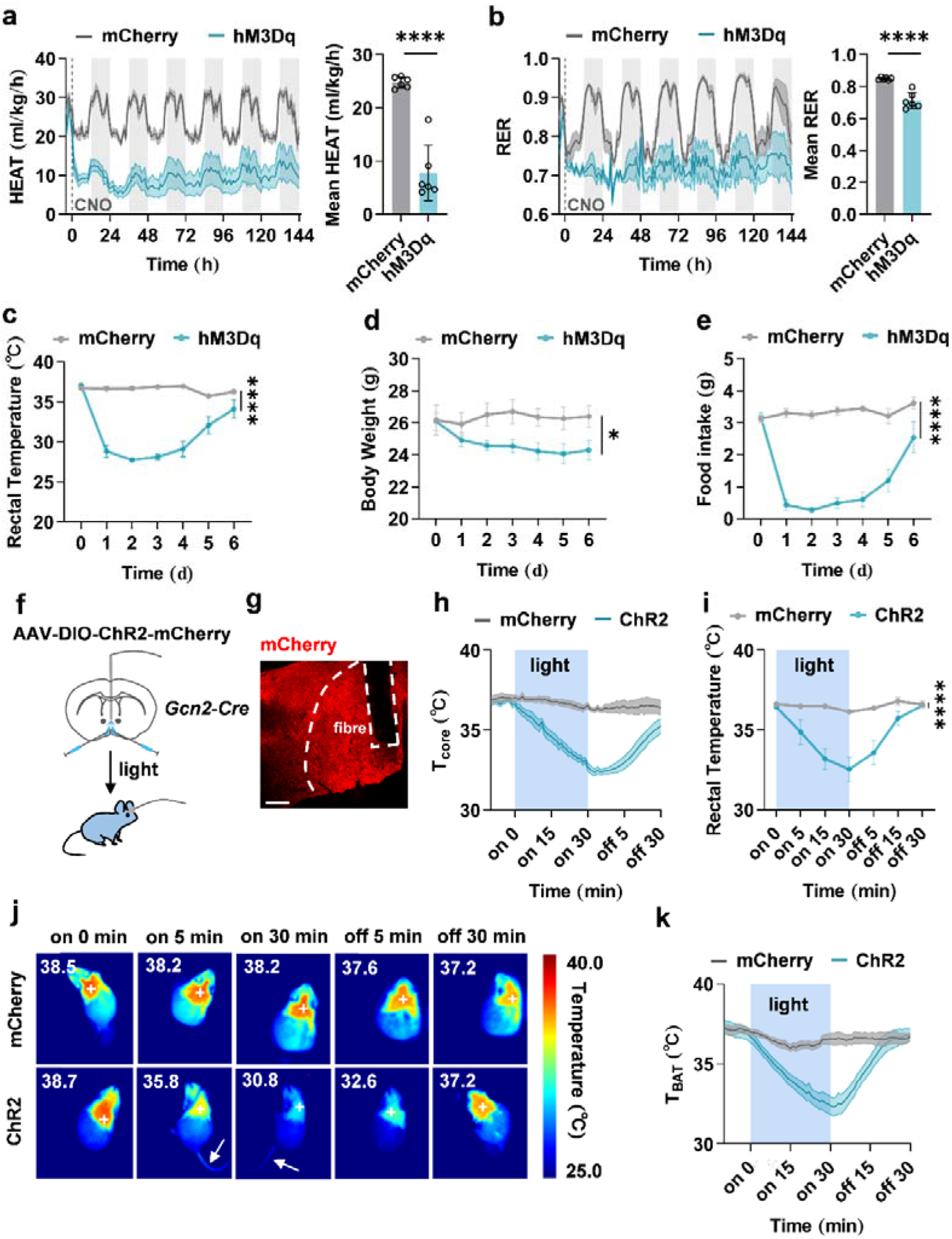
Other effects of activation of G neurons, and consistent effects on thermoregulation of optogenetic activation of G neurons. **a-e,** Other effects of activation of G neurons. **a, b,** Left, energy expenditure (HEAT), and respiratory exchange ratio (RER) before and after CNO administration in control (mCherry) and G-hM3Dq (hM3Dq) mice (n = 6). Right, quantifications of mean HEAT and RER after CNO administration (n = 6, ****p = 1.4e-5, ****p = 6.7e-5, respectively). **c-e,** Rectal temperature, body weight, and food intake following CNO administration (n = 6, ****p = 6e-15, *p = 0.029, ****p = 2.1e-10, respectively). **f-k,** Effects on thermoregulation of optogenetic activation of G neurons. **f,** Experiment strategy. **g,** Representative image of avMPA sections showing ChR2 expression (mCherry) and fiber implantation. **h, i, k,** Core body temperature (T_core_), rectal temperature, and brown adipose tissue surface temperature (T_BAT_) of control (mCherry) and G-ChR2 (ChR2) mice following light stimulation (indicated by blue bars, 20 Hz, 30 min) (n = 5, ****p = 1.5e-7). **j,** Representative infrared thermal images of T_BAT_ at time indicated before, during and after stimulation. Data are represented as mean ± SEM. Two-tailed unpaired Student’s t test in (**a, b**). Two-way RM ANOVA with Geisser-Greenhouse’s correction in (**c, d, e, i**). ****p < 0.0001.

**Extended Data Fig. 5.**
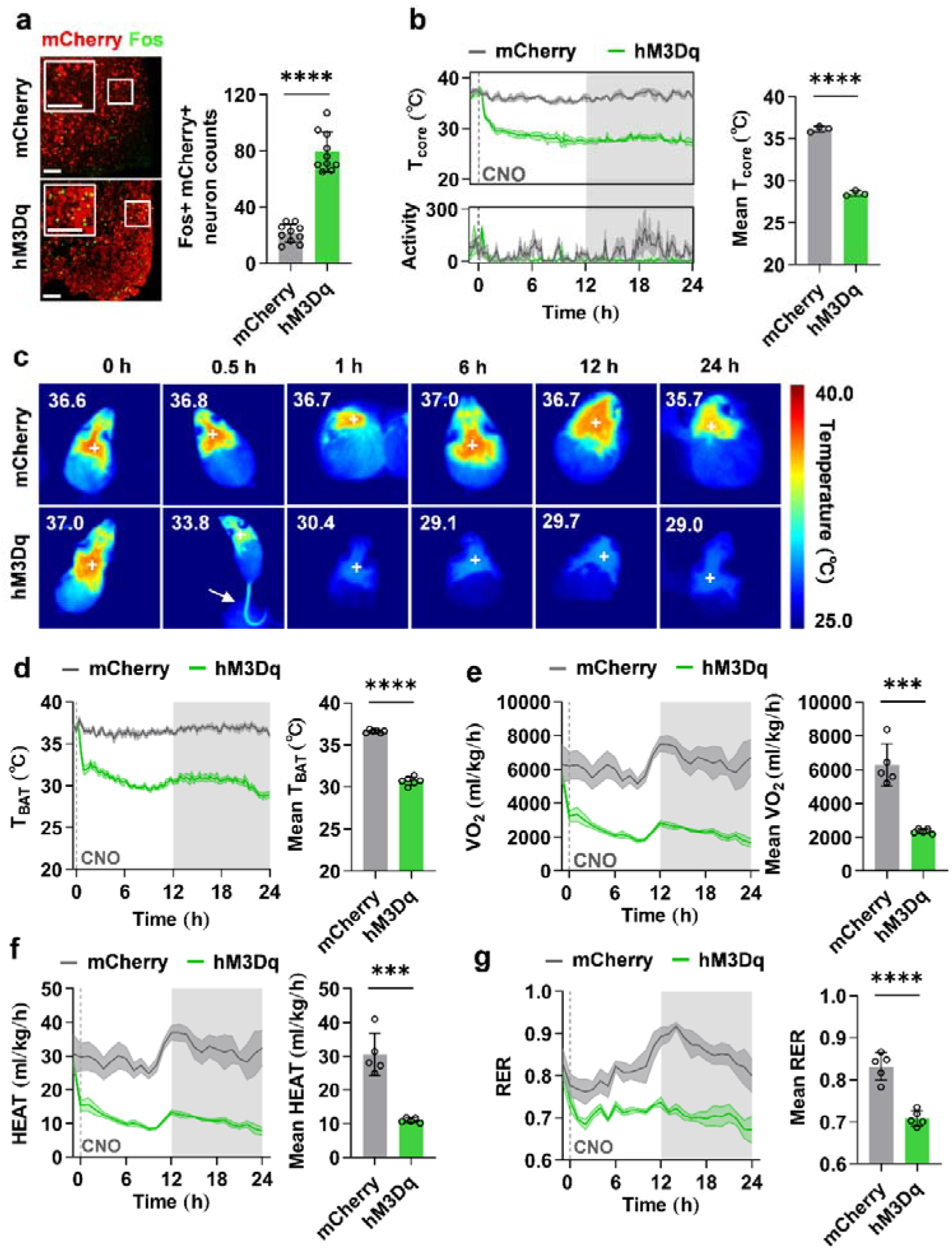
Activation of G neurons also promotes torpor in female mice. **a,** Left, representative immunofluorescence (IF) staining images for Fos (green) in avMPA sections in control (mCherry) and G-hM3Dq (hM3Dq) mice following CNO administration. Right, quantifications of Fos^+^/mCherry^+^ colocalized neuron counts (n = 10, ****p = 7.2e-10). Scale bars: 100 μm for both main panels and close-up. **b,** Left, core body temperature (T_core_) and motor activity (Activity) before and after CNO administration (n = 3). Right, quantifications of minimum T_core_ (n = 3, ****p = 7.6e-6). Gray/white areas indicate 12 hours of darkness/light respectively. Dashed lines indicate the onset of CNO administration. **c,** Representative infrared thermal images of brown adipose tissue surface temperature (T_BAT_) at time indicated before and after CNO administration. Tail temperature increased at 0.5 h (arrow). **d-g,** Left, T_BAT_, metabolic rate (volume of oxygen consumed, VO_2_), energy expenditure (HEAT), and respiratory exchange ratio (RER) before and after CNO administration (n=5). Right, quantifications of mean T_BAT_, VO_2_, HEAT, and RER after CNO administration (n = 5, ****p = 6.3e-11, ***p = 1.2e-4, ***p = 1.3e-4, ****p = 8e-5, respectively). Data are represented as mean ± SEM. Two-tailed unpaired Student’s t test in (**a, b, d, e, f, g**). ***p < 0.001; ****p < 0.0001.

**Extended Data Fig. 6.**
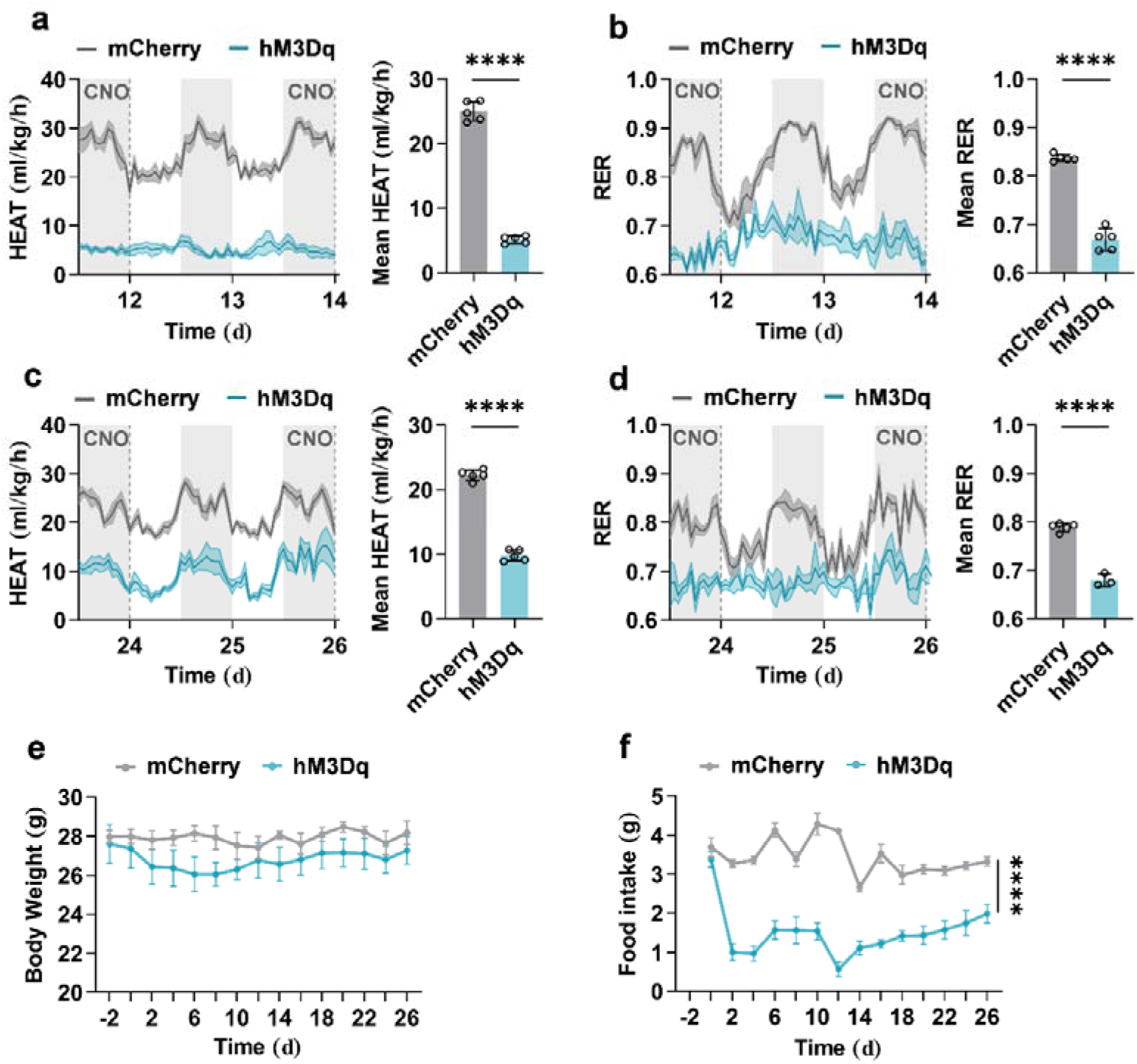
Other phenotypes in G neuron-driven long-term torpor (GLT). **a, b,** Left, energy expenditure (HEAT), and respiratory exchange ratio (RER) of control (mCherry) and G-hM3Dq (hM3Dq) mice during the mid-term (12-14 day) (n = 5). Right, the quantifications of the mean HEAT and RER (n = 5, ****p = 3.7e-9, ****p = 2.6e-7, respectively). **c, d,** Left, the HEAT (n = 5) and the RER (mCherry, n = 5, hM3Dq, n = 3) during the final term (24-26 day). Right, the quantifications of the mean HEAT (n = 5) and the mean RER (mCherry, n = 5, hM3Dq, n = 3, ****p = 1.2e-8, ****p = 6e-6, respectively). **e, f,** Body weight and food intake during GLT (n = 5, p = 0.55, ****p = 4.5e-10, respectively). Data are represented as mean ± SEM. Two-way RM ANOVA with Geisser-Greenhouse’s correction in (**e, f**). ****p < 0.0001.

**Extended Data Fig. 7.**
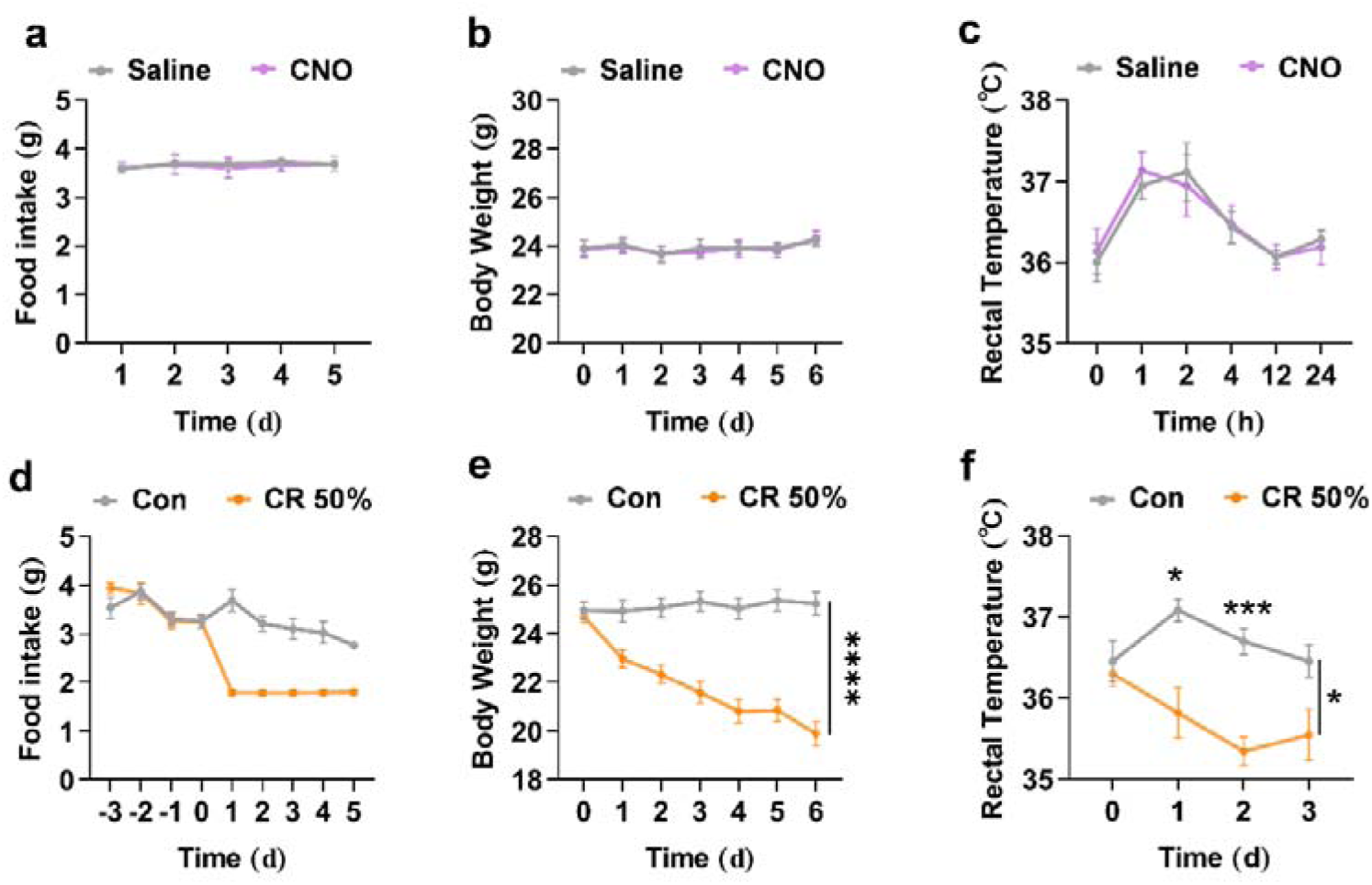
Caloric restriction and drug administration alone are insufficient to cause long-term torpor. **a-c,** Mice were subjected to saline/CNO administration every day for 5 d. Food intake (**a**), body weight (**b**) during the 5 d and the rectal temperature (**c**) in 24 h after administration showed no significant change (n = 6, p = 1, p = 0.93, p = 0.97, respectively). **d-f,** Caloric restriction (CR) was insufficient to cause long-term torpor. **d,** Food intake of mice ad libitum fed (Con) and mice subjected to 50 % of their average daily food intake (CR 50%, n = 7). **c, d,** Body weight and rectal temperature (****p < 1e-15) of Con and CR 50 % mice (n = 7, *p = 0.03 between lines, p = 1, *p = 0.021, ***p = 4.1e-4, p = 0.14, respectively at indicated time points). A slight decrease in rectal temperature was observed. Data are represented as mean ± SEM. Two-way RM ANOVA with Geisser-Greenhouse’s correction in (**b, d, e, f**). Two-way RM ANOVA with Geisser-Greenhouse’s correction followed by post hoc unpaired t-test with Bonferroni’s correction in (**c**). *p < 0.05; ****p < 0.0001.

**Extended Data Fig. 8.**
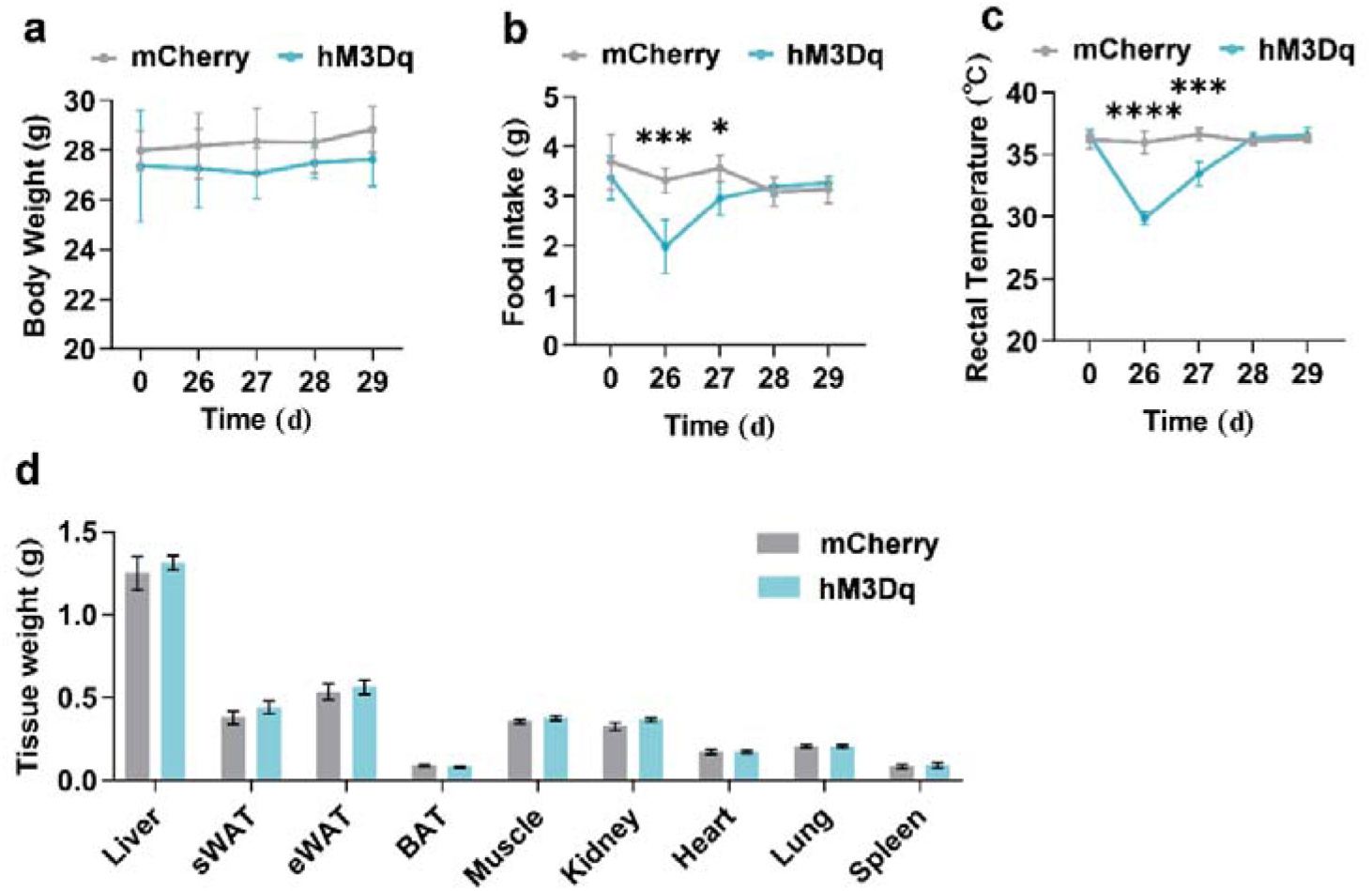
Other phenotypes following G neuron-driven long-term torpor (GLT). **a-c,** Body weight, food intake and rectal temperature following GLT in control (mCherry) and G-hM3Dq mice (hM3Dq, n = 5, p = 0.95, p = 0.076, p = 0.14, p = 0.22, p = 0.095 in **a,** p = 0.34, ***p = 9.9e-4, *p = 0.012, p = 0.49, p = 0.38 in **b,** p = 0.54, ****p = 1.1e-6, ***p = 1.7e-4, p = 0.29, p = 0.35 in **c,** respectively). **c,** Tissues weight of liver, subcutaneous white adipose tissue (sWAT), epididymal white adipose tissue (eWAT), brown adipose tissue (BAT), muscle, kidney, heart, lung and spleen. (mCherry, n = 8, hM3Dq, n = 9, p = 0.56, p = 0.28, p = 0.66, p = 0.20, p = 0.28, p = 0.15, p = 0.96, p = 0.96, p = 0.68, respectively). Data are represented as mean ± SEM. Two-tailed unpaired Student’s t test in (**a-c**). *p < 0.05; ***p < 0.001; ****p < 0.0001.

**Extended Data Fig. 9.**
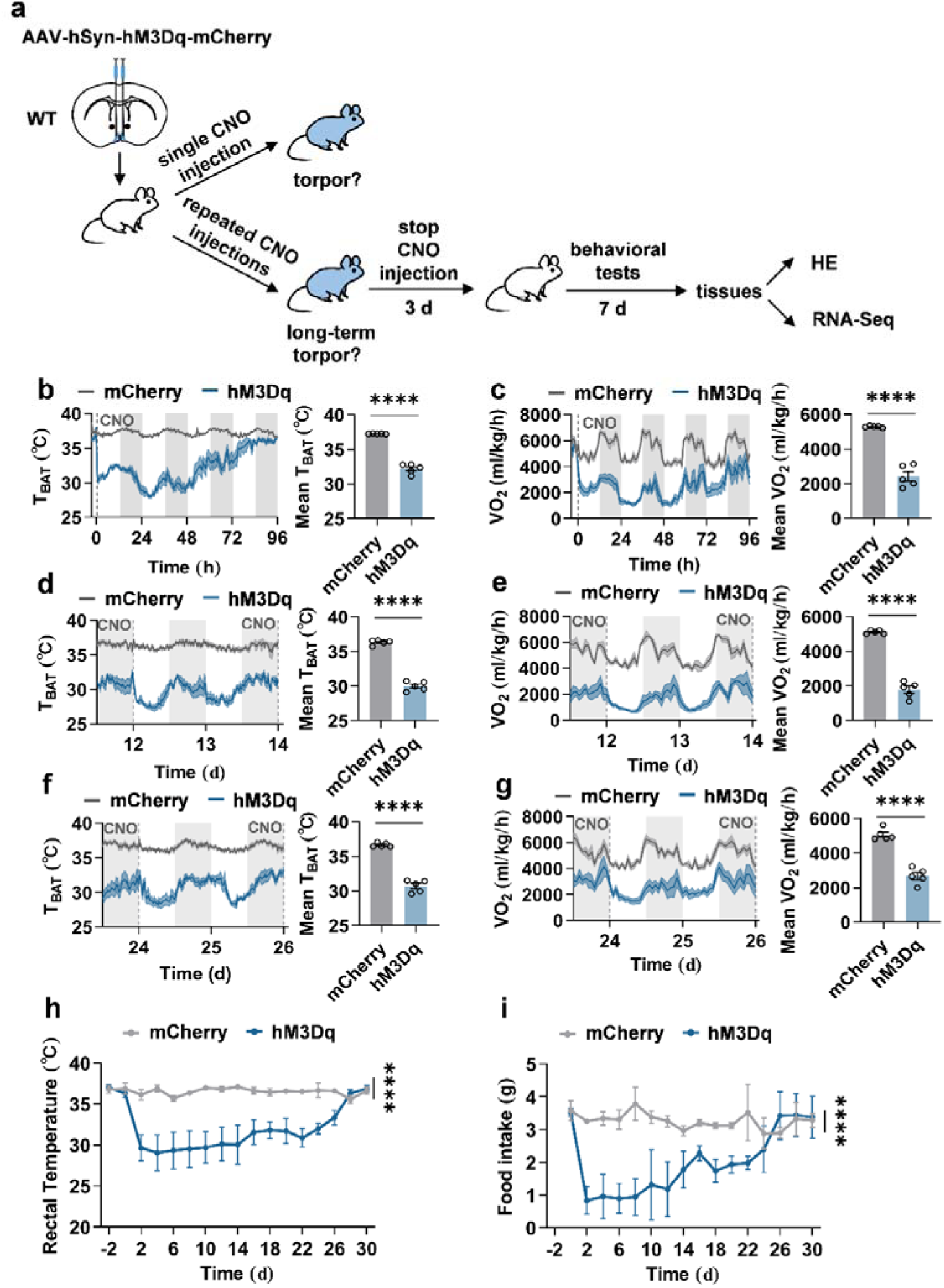
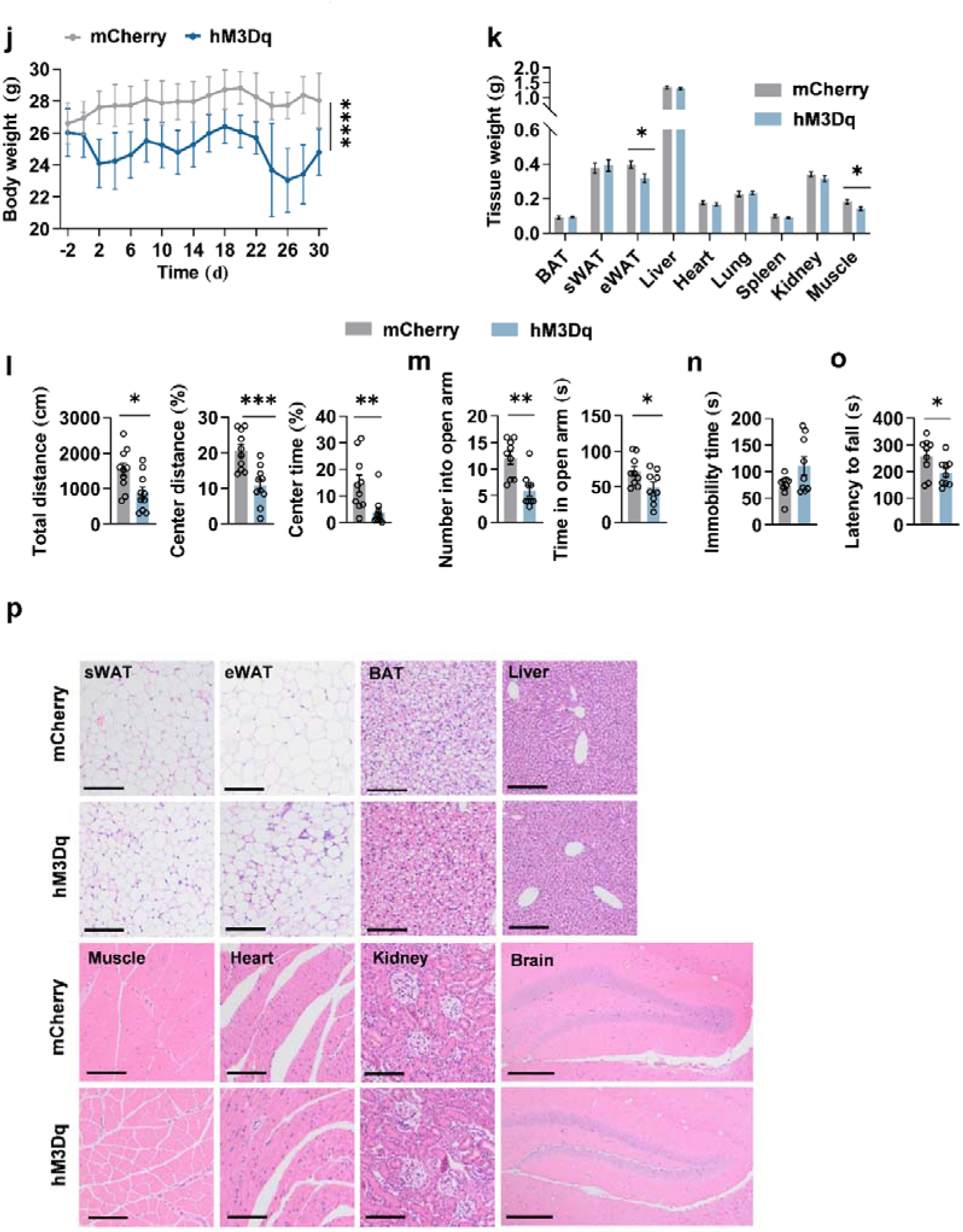
Induction of pan-neuron-driven long-term torpor (PLT) and secondary pathological changes following it. **a,** Experimental strategy for single or repeated activation of pan-neurons in avMPA. Hematoxylin and eosin (HE). **b, c,** Effects of single CNO administration in control (mCherry) and pan-neurons-hM3Dq (hM3Dq) mice. Left, brown adipose tissue surface temperature (T_BAT_) and metabolic rate (volume of oxygen consumed, VO_2_) before and after CNO administration (n = 5). Right, quantifications of mean T_BAT_ and mean VO_2_ after CNO administration (n = 5, ****p = 2.4e-7, ****p = 7.0e-6, respectively). Gray/white areas indicate 12 hours of darkness/light, respectively. Dashed lines indicate the onset of CNO administration. **d-g,** Phenotypes measurement in PLT. **d, e,** Left, T_BAT_ and VO_2_ during mid-term (12–14 day). Right, quantifications of the mean T_BAT_ and the mean VO_2_ (n = 5, ****p = 6.5e-8, ****p = 2.4e-7, respectively). **f, g,** Left, T_BAT_ and VO_2_ during final term (24–26 day). Right, the quantifications of the mean T_BAT_ and VO_2_ (n = 5, ****p = 5.7e-7, ****p = 1.4e-5, respectively). **h-j,** Rectal temperature, food intake and body weight during and following PTL (n = 5, ****p < 1e-15, ****p < 1e-15 ****, p = 4.2e-8, respectively). **k,** Tissues weight of liver, subcutaneous white adipose tissue (sWAT), epididymal white adipose tissue (eWAT), brown adipose tissue (BAT), muscle, kidney, heart, lung and spleen. (n = 7, p = 0.81, p = 0.73, *p = 0.032, p = 0.46, p = 0.54, p = 0.70, p = 0.38, p = 0.26, *p = 0.021, respectively). **l-n,** Behavior deficits and tissues damages observed following PLT. **l,** Total travel distance (Total distance), percentage of distance in center area (Central distance), and percentage of time spent in center (Central time) of open filed test (OFT) following the induction in mCherry and hM3Dq mice (n = 10, *p = 0.017, ***p = 6.9e-4, **p = 8.8e-3, respectively). **m,** Number of entries into the open arms (Number into open arm) and time spent in the open arms (Time in open arm) of elevated plus maze (EPM) (n = 9, **p = 1.5e-3, *p = 0.041, respectively). **n,** Immobility time of tail suspension test (TST) (n = 9, p = 0.065). **o,** Latency to fall of rotarod test (RR) (n = 9, *p = 0.045). **p,** Representative images of HE staining of subcutaneous white adipose tissue (sWAT), epididymal white adipose tissue (eWAT), brown adipose tissue (BAT), liver, muscle, heart, kidney and brain. Scale bars: 200 μm for sWAT, eWAT, BAT, muscle, heart, kidney and brain; 400 μm for liver. Data are represented as mean ± SEM. Two-tailed unpaired Student’s t test in (**b-g, l-o**). Two-way RM ANOVA with Geisser-Greenhouse’s correction followed by post hoc unpaired t-test with Bonferroni’s correction in (**h-j**). *p < 0.05; **p < 0.01; ***p < 0.001; ****p < 0.0001.

**Extended Data Fig. 10.**
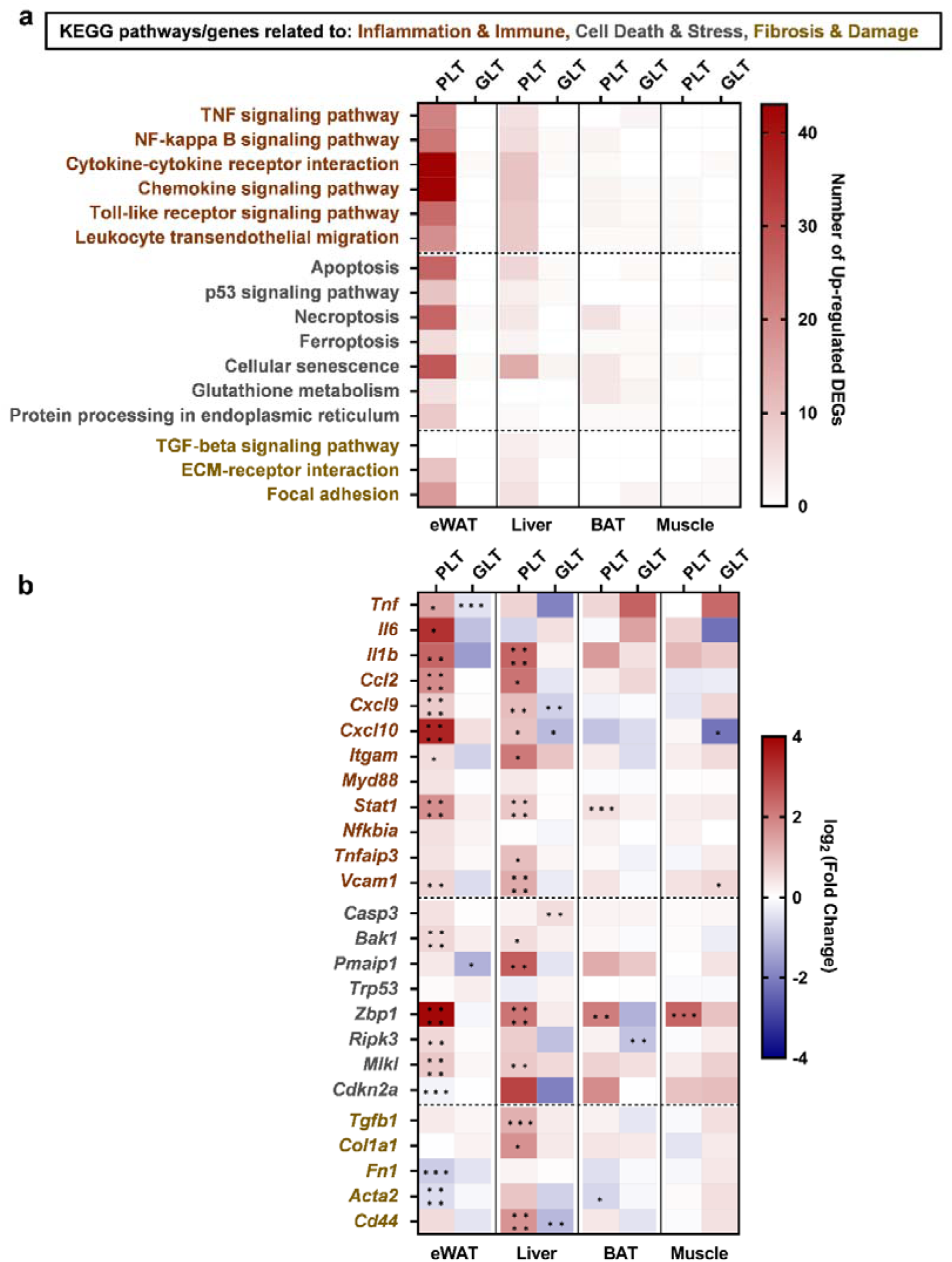
Transcriptomic profiling reveals mitigated systemic toxicity and tissue damage following G neuron-driven torpor (GLT). **a,** Heatmap showing the number of up-regulated differentially expressed genes (DEGs) enriched in selected KEGG pathways associated with tissue damages and pathological responses in epididymal white adipose tissue (eWAT), liver, brown adipose tissue (BAT), muscle following GLT and pan-neuron-driven long-term torpor (PLT). The pathways are categorized into three main functional blocks: Inflammation & Immune (brown), Cell Death & Stress (grey), and Fibrosis & Damage (gold). The color gradient (from white to dark red, 0 to 43) represents the absolute count of up-regulated DEGs within each pathway across four tissues for the indicated treatment groups (PLT and GLT) (n = 3). **b,** Heatmap showing the transcriptional expression levels of representative marker genes corresponding to the three functional categories defined in (**a**). The color scale (from dark blue to dark red, -4 to 4) indicates the log_2_ (Fold Change) of gene expression in the PLT or GLT groups relative to their respective independent control groups (n = 3). Red indicates up-regulation (harmful effects), while blue indicates down-regulation (mitigation or protective response). Data in (**b**) are represented as *log_2_* (Fold Change). Differential expression analysis was performed using the Wald test (DESeq2) in (**b**). *p-adj < 0.05; **p-adj < 0.01; ***p-adj < 0.001; ****p-adj < 0.0001.

**Extended Data Fig. 11.**
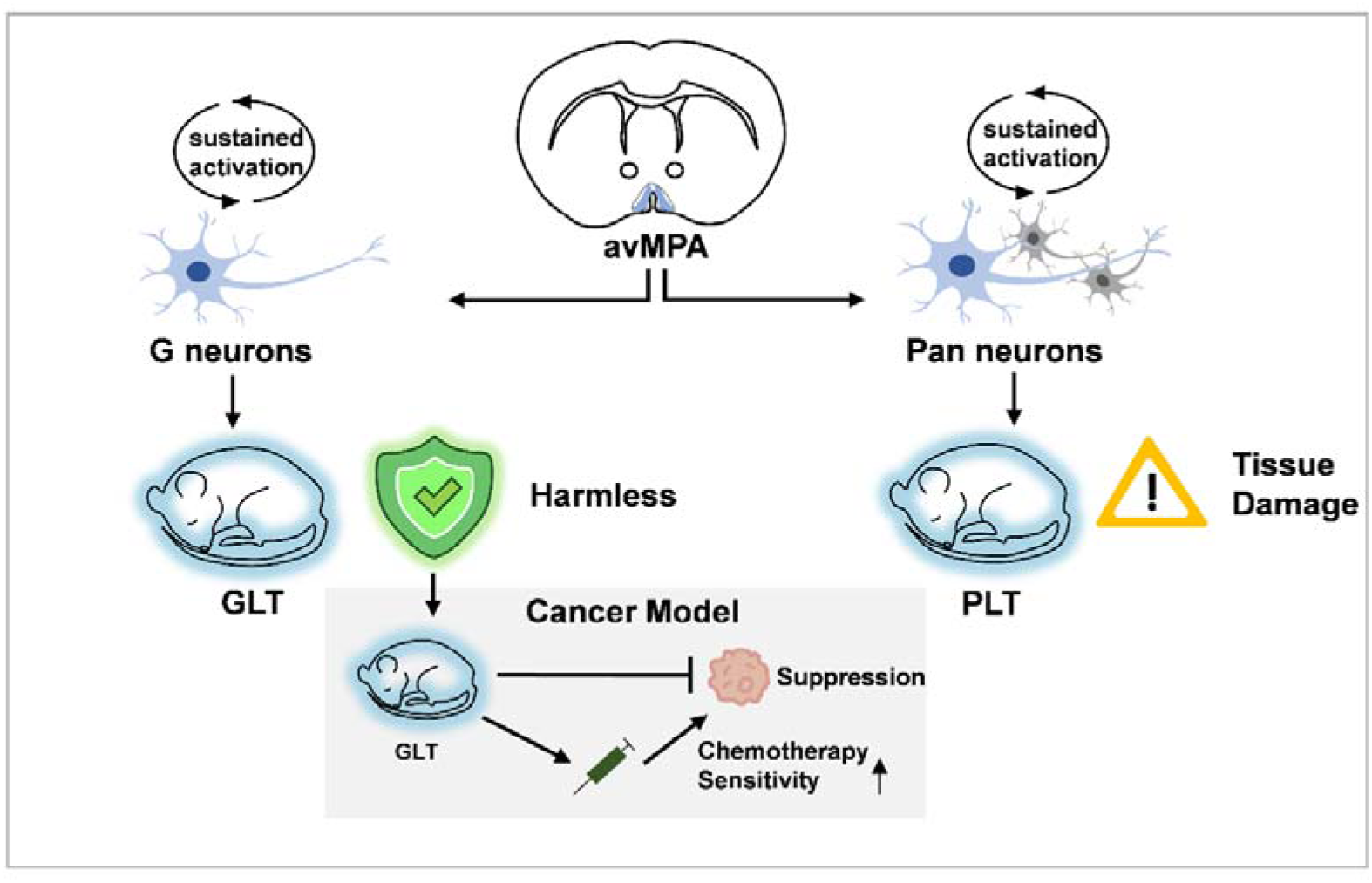
Working model of safely inducing G neuron-driven long-term torpor (GLT). Sustained activation of G neurons safely induces a prolonged torpor in mice. In contrast, sustained pan-neuronal activation in the same brain region (avMPA) results in a long-term torpor (PLT) with tissue damage. GLT can both directly suppress tumor growth and enhances the chemotherapy sensitivity in a mouse cancer model.

